# DNA aptamers that modulate biological activity of model neurons

**DOI:** 10.1101/2024.04.08.588576

**Authors:** Jenelle Rolli, Keenan Pearson, Brandon Wilbanks, Sybil C. L. Hrstka, Arthur E. Warrington, Nathan P. Staff, L. James Maher

## Abstract

There is an urgent need for agents that promote health and regeneration of cells and tissues, specifically to treat diseases of the aging nervous system. Age-associated nervous system degeneration and various diseases are driven by many different biochemical stresses, often making it difficult to target any one disease cause. Our laboratory has previously identified DNA aptamers with apparent regenerative properties in murine models of multiple sclerosis by selecting aptamers that bind oligodendrocyte membrane preparations. Here, we screened vast libraries of molecules (∼10^14^ unique DNAs) for the ability to bind cultured human SH-SY5Y neuroblastoma cells as model neurons to demonstrate the feasibility of identifying biologically active aptamers by cycles of cell selection. Many of these DNA aptamers bind undifferentiated and differentiated cultured SH-SY5Y cells. Several of these aptamers modulate the biological activity of SH-SY5Y cells upon treatment in culture.

## Introduction

The etiologies of neurodegenerative diseases vary widely even within a single disease category, and many of these diseases share genetic links and patterns of molecular pathogenesis. This variability in etiology raises the possibility of generic disease treatment by stimulating cell health, resilience, and regeneration, ideally countering neurodegeneration stemming from a wide range of causes. Approaches that seek to promote cell regeneration have been undertaken utilizing stem cells, antibodies, and nucleic acid aptamers. There are several stem cell-based therapies currently undergoing clinical trials (NCT03280056, NCT03268603, NCT03482050). These therapies have the goal of neuroprotection through paracrine effects, rather than the replacement of lost neurons. One approach relies on autologous mesenchymal stromal cells (MSCs) for their ability to secrete neurotrophic factors and modulate the immune system.^1–3^ Various strategies are being implemented to increase neurotrophic factor secretion by such MSCs, including conditioned media and gene editing to drive overexpression.^4, 5^ Another approach uses neuroglial lineage precursors.^6, 7^ However, the allogeneic nature of this formulation requires concomitant immunosuppression.

Certain natural human IgM antibodies have been identified for their ability to bind neurons and elicit regenerative responses.^8–17^ A recombinant form of one such human IgM, rHIgM22, was shown to promote central nervous system (CNS) remyelination in models of multiple sclerosis (MS),^9–11, 14, 17, 18^ a disease in which the myelin sheath coating neuronal axons is damaged. Based on the results reported for rHIgM22, our laboratory developed DNA aptamers LJM-3064 and LJM-5708 by cycles of selection for binding to rodent CNS myelin suspensions.^18^ We then studied their pharmacodynamics and delivery to the CNS of mice after intraperitoneal (i.p.) injection of these DNA aptamers.^19^ When biotinylated and multimerized around a streptavidin core, formulations of these aptamers showed features reminiscent of rHIgM22, promoting myelin regeneration and functional improvement in a Theiler’s Murine Encephalomyelitis Virus (TMEV) mouse model of progressive MS.^10, 18–21^ An independent study showed efficacy of a different formulation of this DNA aptamer in an experimental autoimmune encephalomyelitis (EAE) mouse model of MS.^22^ This work demonstrates the potential for selection of biologically active DNA aptamers by affinity selection against disease-relevant targets.

Another neuron-binding human antibody reported to elicit regenerative responses is rHIgM12. This molecule has been shown to increase neuron attachment in culture and promote neurite outgrowth.^12, 16^ rHIgM12 was further evaluated for in vivo efficacy in two SOD1 mouse models of Amyotrophic Lateral Sclerosis (ALS).^15^ A single dose of rHIgM12 was reported to increase survival, reduce symptom progression, reduce myelin whorls (indicative of degenerating axons), and increase the number of spinal cord anterior horn cells (indicating a reduction in motor neuron degeneration) in mice.^15^ Such results might translate into increased human ALS patient survival.

Inspired by these studies hinting at the potential for neuroprotective and neuroregenerative molecules, we used unbiased in vitro selection to identify DNA aptamers that specifically bind SH-SY5Y neuroblastoma cells, commonly used in neurology research as model neurons.^23–25^ This strategy provided a library of molecules that can be screened for therapeutic properties. Such aptamers might serve as targeted delivery agents for therapeutic molecules, or more interestingly, cellular binding might itself trigger regenerative responses in the context of neurodegenerative diseases, nerve injuries, and nerve grafts. With cell culture assays of growth, morphology, and elicited transcriptional responses, we describe several DNA aptamers that modulate biological activity in SH-SY5Y cells.

## Results

### Aptamer selection

We set out to determine if DNA aptamers can be identified with the ability to elicit potentially useful biological responses in neurons. Our strategy was to identify cell-binding DNA aptamers by in vitro selection, characterize and quantitate binding specificity, and then screen for biological activity. To do this, cultured adherent SH-SY5Y cells were first used as a convenient neuron model for positive selection and non-neuronal cells as negative selection targets to identify SH-SY5Y-specific DNA aptamers using the three similar approaches described in Materials and Methods (Fig. 1A). The progress of each selection was monitored by subjecting recovered aptamer libraries to qPCR.^26^ In general, library recovery increased across selection rounds (Fig. S1). The original naïve library (denoted selection round 0) and libraries from various selection rounds were subjected to Next Generation Sequencing. Top sequences were identified based on their enrichment over rounds of selection, indicated by their abundance (Fig. 1B-D). The proportion of non-unique sequences increased over the course of each selection (Fig. S2).

**Fig. 1.**
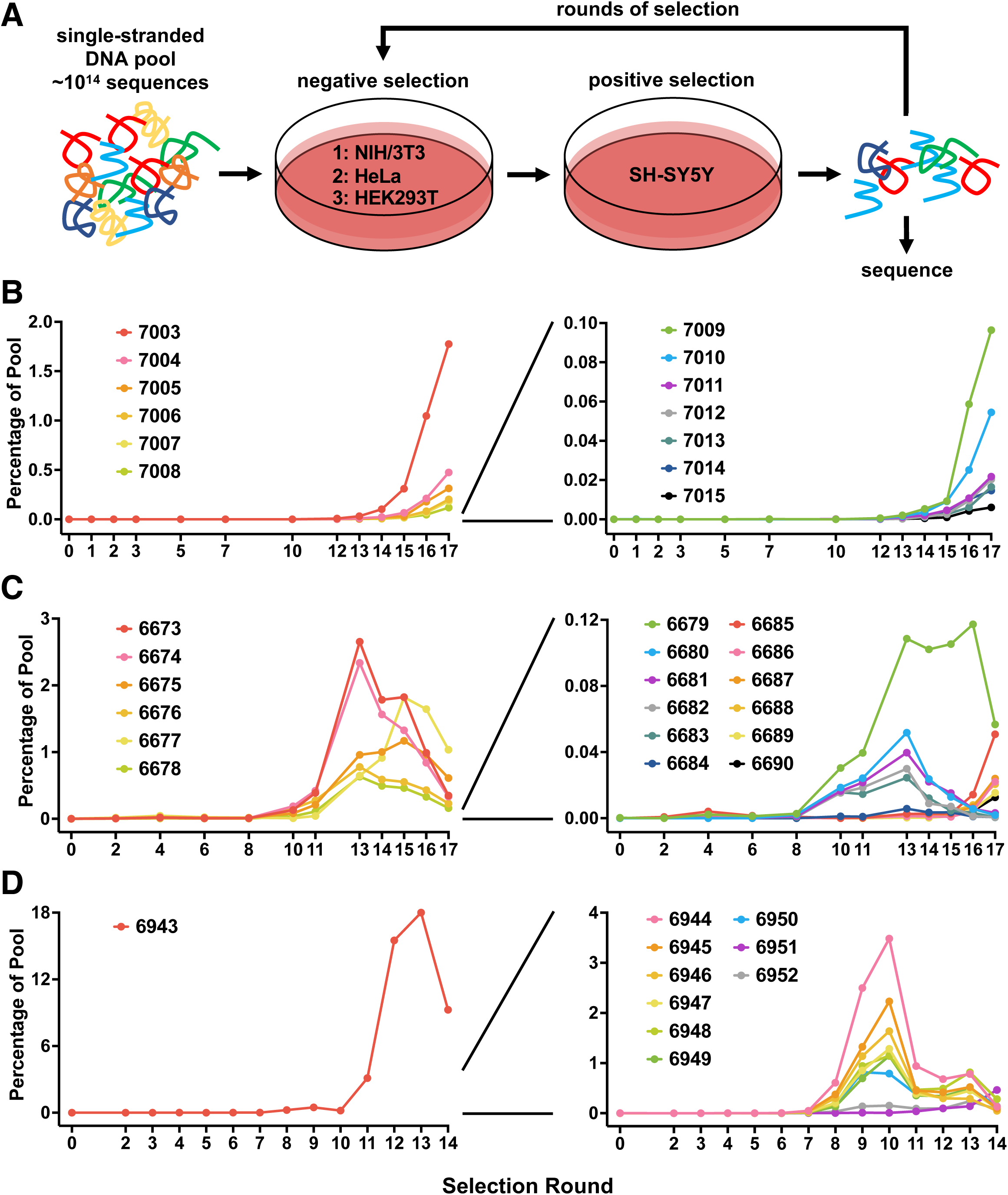
Selection of DNA aptamers that bind SH-SY5Y cells. A) Schematic of selection procedure showing negative selection cell lines used for selections 1, 2, and 3. B-D) Prevalence of the indicated candidate from deep sequencing data is shown across rounds of selections 1-3, respectively. Panels on right expand the y-axis for clarity.

We chose 41 unrelated sequences from the top candidates identified in each of the three similar selections (Fig. 1A) using AptaSUITE.^27^ These candidates were synthesized individually with 3’-biotin tags and screened for the ability to specifically bind model neurons (Fig. 1B-D and Table S1). We also chose negative control sequences 6699 and 6953 that were present in the original naïve selection libraries but not in any of the rounds of selection (Table S1). The predicted secondary structures of all candidates were generated using Mfold and the predicted free energies of the negative control oligonucleotides were within the range of those for the candidate aptamers.^28, 29^

### Aptamer binding

We assessed the binding of all candidate aptamer sequences and controls to SH-SY5Y cells by image analysis. SH-SY5Y cells were seeded and grown overnight to the same density as in the selection (Fig. 2 inset). Aptamer staining was performed and quantified as described in Materials and Methods. Three previously described DNA aptamers were included along with the candidate aptamers and controls described above. First, we included a previously published aptamer from our laboratory, 3064, which had been selected against rodent CNS myelin suspension.^18^ Another aptamer, yly12, was reportedly selected against neurites isolated from differentiated SH-SY5Y cells, but also bound undifferentiated SH-SY5Y cells. This aptamer reportedly binds L1CAM.^30^ The third aptamer, Apt3, was reported to bind to poly-sialic acid and was of interest because neuroregenerative antibody rHIgM12 reportedly binds to poly-sialic acid modified NCAM.^13, 31^ All candidates were synthesized with 3’ terminal biotin tags allowing detection of cell-specific binding by fluorescently labeled streptavidin following incubation on cultured cells, washing to remove unbound molecules, and formaldehyde fixation. Twenty-four candidate aptamers, including all those previously published, exhibited statistically significant staining of SH-SY5Y cells compared to both negative control oligonucleotides (Fig. 2). We found that pre-conjugation of biotinylated aptamers with fluorescent streptavidin resulted in optimal staining, so we optimized the ratio of aptamer to streptavidin to minimize free aptamer and maximize higher order multimers (Fig. S3). The staining of several aptamers was confirmed by high-resolution confocal microscopy using the optimized pre-conjugation method (Fig. S4).

**Fig. 2.**
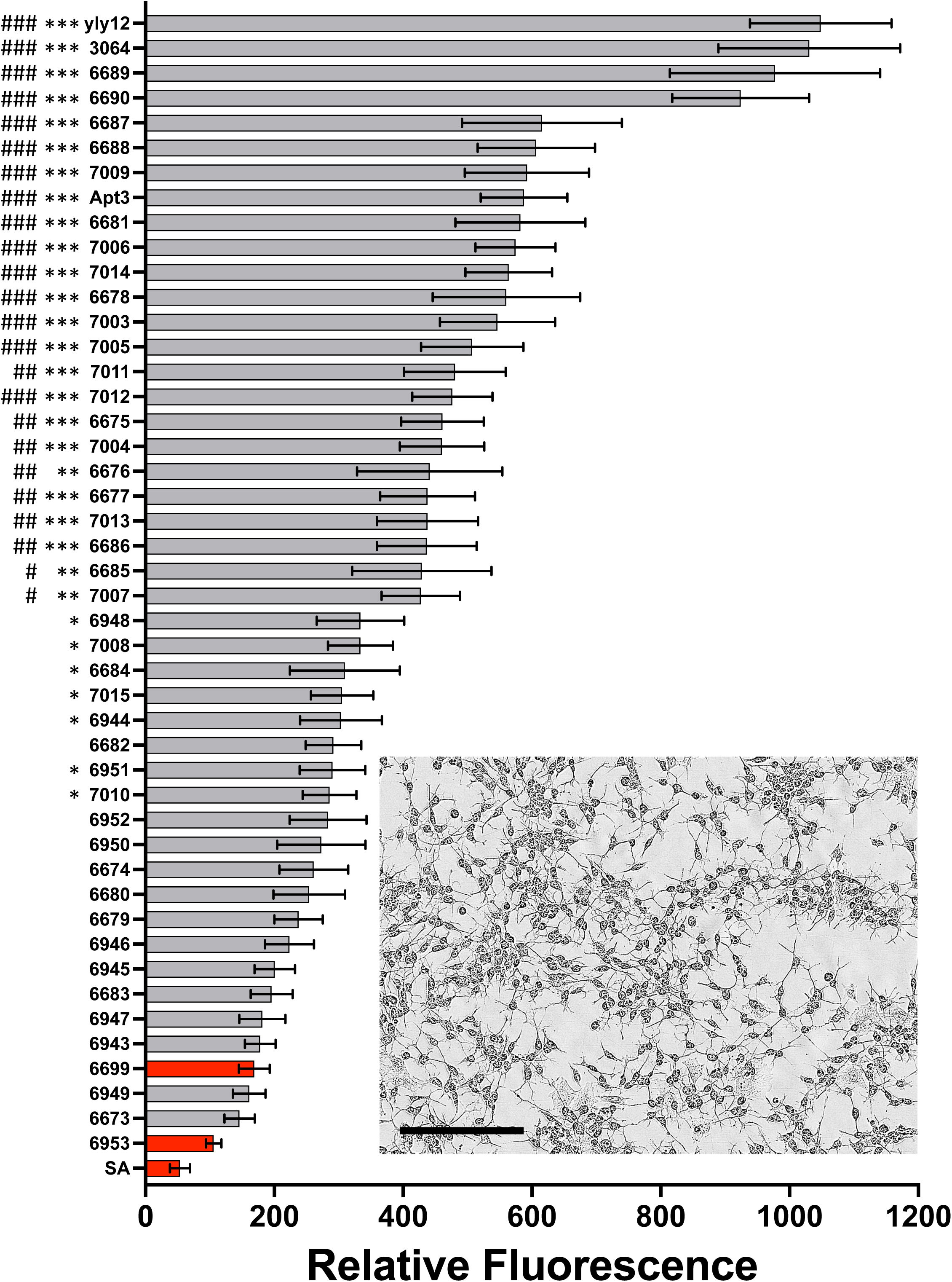
Screening candidate aptamers by assessing binding to SH-SY5Y cells. Phase contrast image to show cell morphology of proliferating SH-SY5Y cells at 20× magnification, scale bar is 200 µm (inset). Binding quantification for candidates (7003 – 7015, 6673 – 6690, and 6943 – 6952), other published aptamers (3064, 6214, and 6565), and negative controls (6699, 6953, and SA; bars shown in red). Quantification of staining was calculated as NIR signal intensity per image field normalized to confluence (as described in Materials and Methods). Statistical significance is shown for comparison with 6699 (#: p<0.05, ##: p<0.01, ###: p<0.001) and 6953 (*: p<0.05, **: p<0.01, ***: p<0.001) by one-way ANOVA. Data are represented as mean ± SEM for multiple image fields (n=34-78). Image analysis was performed with IncuCyte SX5.

We sought to optimize certain aptamers by truncation. We used Mfold to predict the secondary structures of ten aptamers.^28^ Truncations were then designed to remove apparently unstructured terminal regions. In most cases, the original aptamers performed better or equally relative to truncates, except for 6686 and 6952 where the truncated versions significantly improved staining over their full-length counterparts (Fig. 3).

**Fig. 3.**
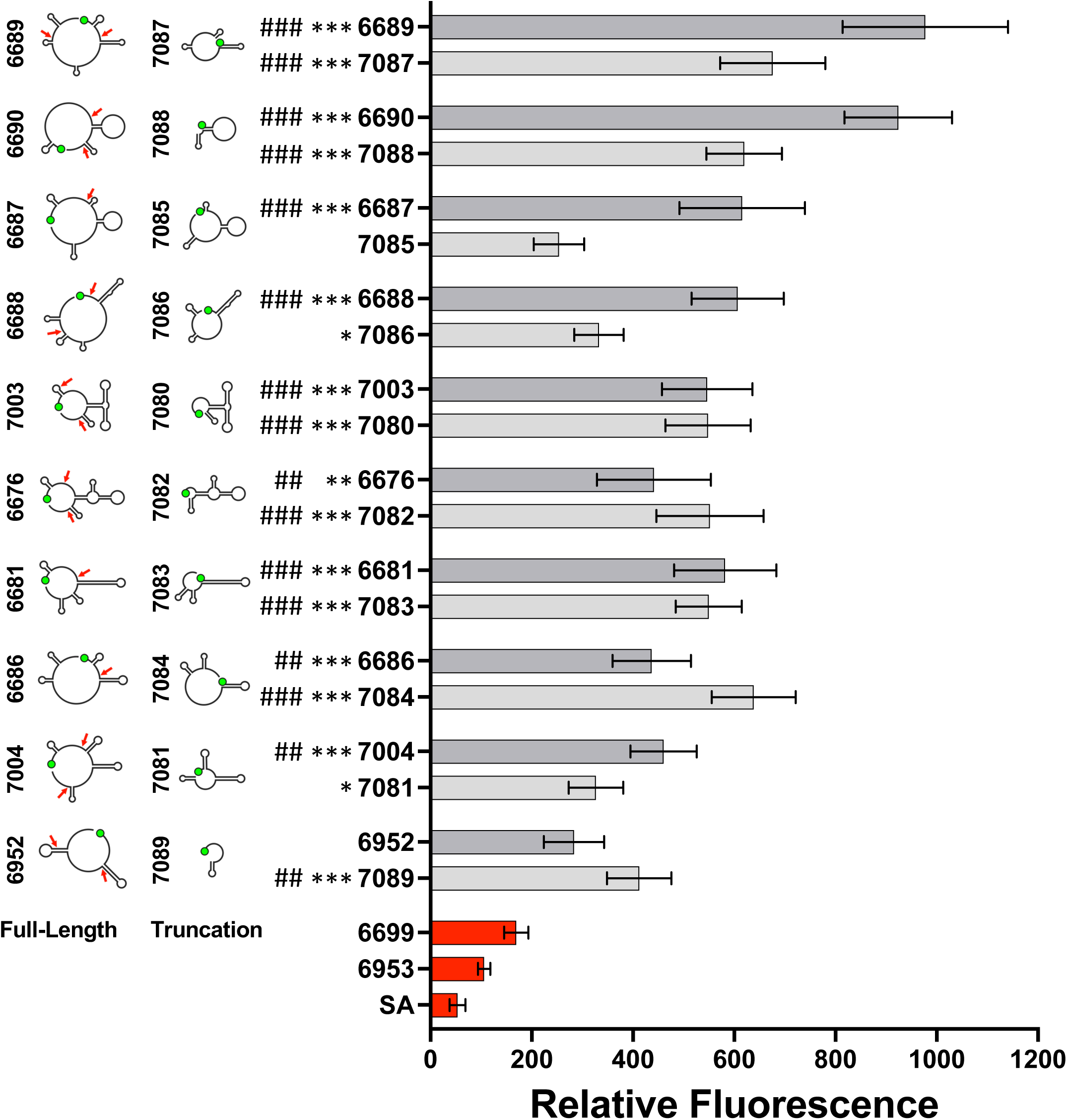
Screening truncated aptamers by assessing binding to SH-SY5Y cells. Schematic representations of predicted secondary structures for full-length aptamers and truncations determined using mFold. Green circle indicates the 5’ terminus. Red arrows indicate truncation positions within full-length sequences. Binding quantification of full-length aptamers are shown in the top of each bar grouping (dark gray), and quantification of truncations are shown in the bottom of each bar grouping (light gray). Statistical significance is shown for comparison with 6699 (#: p<0.05, ##: p<0.01, ###: p<0.001) and 6953 (*: p<0.05, **: p<0.01, ***: p<0.001) by one-way ANOVA. Data are represented as mean ± SEM for multiple image fields. Data for full-length aptamers and negative controls are the same as shown in Fig. 2.

We next questioned whether the aptamers selected against undifferentiated SH-SY5Y cells would also bind to their differentiated counterparts. We consider differentiated SH-SY5Y cells to represent a more mature neuron model. A standard protocol for the differentiation of SH-SY5Y cells was used and aptamer binding was assessed at the end of the 11-day induction.^32^ Differentiated SH-SY5Y cells exhibited an obvious increase in the number and length of neurite-like processes potentially establishing contacts between cells (Fig. S5 and Fig. 4 inset). Seventeen of the aptamers that bound undifferentiated cells also exhibited statistically significant staining of differentiated SH-SY5Y cells compared to both negative control oligonucleotides (Fig. 4). The staining of several aptamers was confirmed by high-resolution confocal microscopy using the optimized pre-conjugation method (Fig. S6).

**Fig 4.**
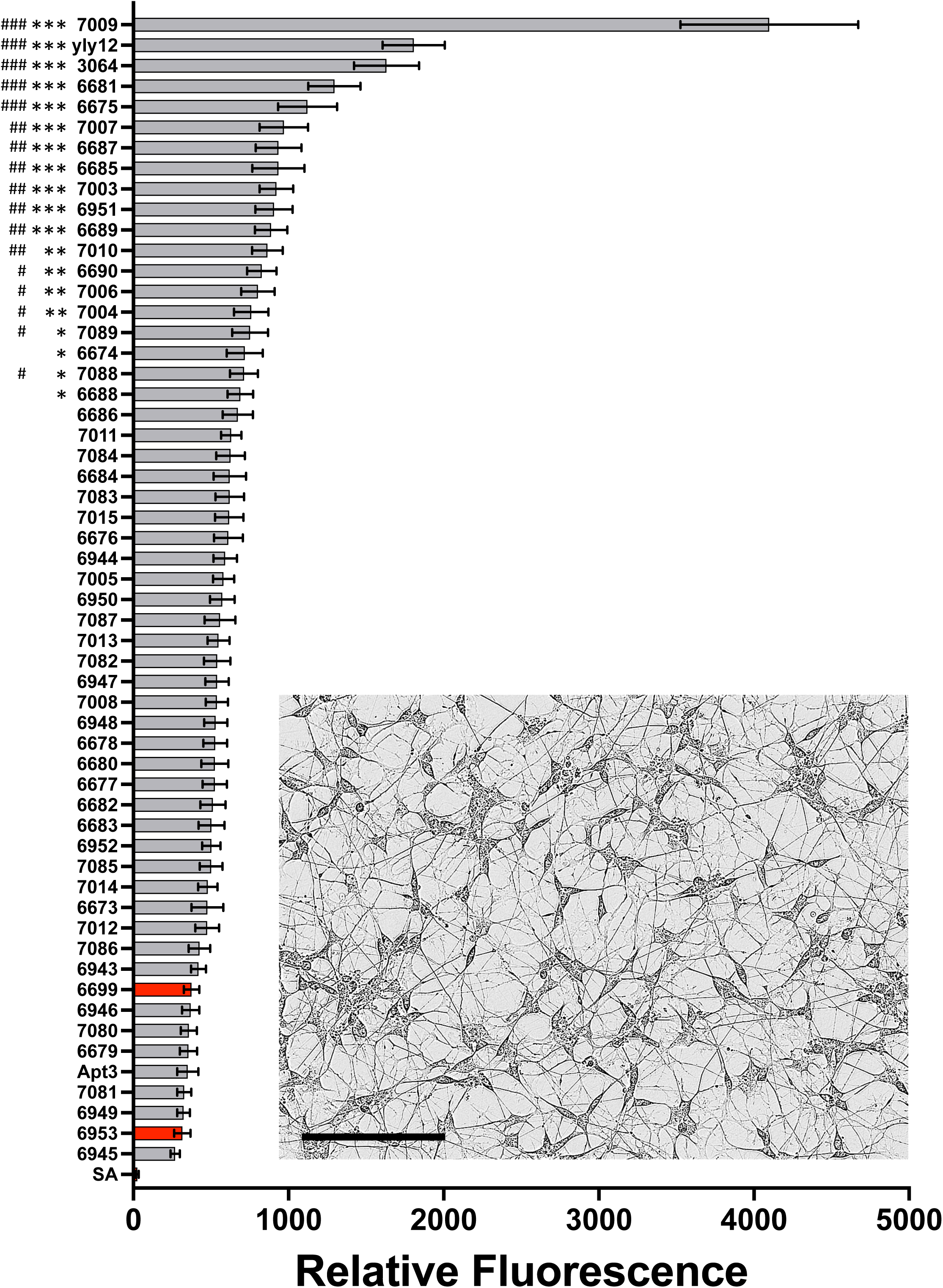
Screening candidate aptamers by assessing binding to differentiated SH-SY5Y cells. Phase contrast image to show morphology of differentiated SH-SY5Y cells at 20× magnification, scale bar is 200 µm (inset). Binding quantification of candidates (7003 – 7015, 6673 – 6690, and 6943 – 6952), other published aptamers (3064, 6214, and 6565), and negative controls (6699, 6953, and SA; bars shown in red). Quantification of staining was calculated as NIR signal intensity per image field normalized to confluence (as described in Materials and Methods). Statistical significance is shown for comparison with 6699 (#: p<0.05, ##: p<0.01, ###: p<0.001) and 6953 (*: p<0.05, **: p<0.01, ***: p<0.001) by one-way ANOVA. Data are represented as mean ± SEM for multiple image fields (n=32-76). Image analysis was performed with IncuCyte SX5.

### Biological assays

After identifying aptamers that bound to target cells, we performed a colony formation assay with the SH-SY5Y neuroblastoma cells to observe any long-term effects of aptamer treatment on cell growth. In this assay, we interpret colony formation as a primary readout of biological activity, including positive or negative growth effects or inducing differentiation. We hypothesize that aptamers which inhibit colony formation may do so by toxic or differentiation effects; however, colony formation counts do not differentiate such responses. The focus of this assay was to narrow our list of candidates with potential biological activity. We observed after 14 days of treatment with 200 nM aptamer that seven candidates altered with statistical significance the ability of SH-SY5Y cells to form colonies. Two candidates increased colony formation and five decreased colony formation relative to a negative control (Fig. 5A and Fig. S7). Because pre-conjugation of biotinylated aptamers with streptavidin had appeared to improve staining results, we tested whether multimerizing biotinylated aptamers with streptavidin would modify the biological response in the colony formation assay. The biological effect of yly12, the aptamer which most effectively inhibited colony formation, was abrogated by multimerization with streptavidin (Fig. S8). We next sought to optimize the concentration of aptamer for maximum effect on colony formation. A dose-response study showed a minor growth inhibitory effect due to the negative control observed at 500 nM, but not at 350 nM (Fig. S9). We therefore used 350 nM aptamer for further testing.

**Fig 5.**
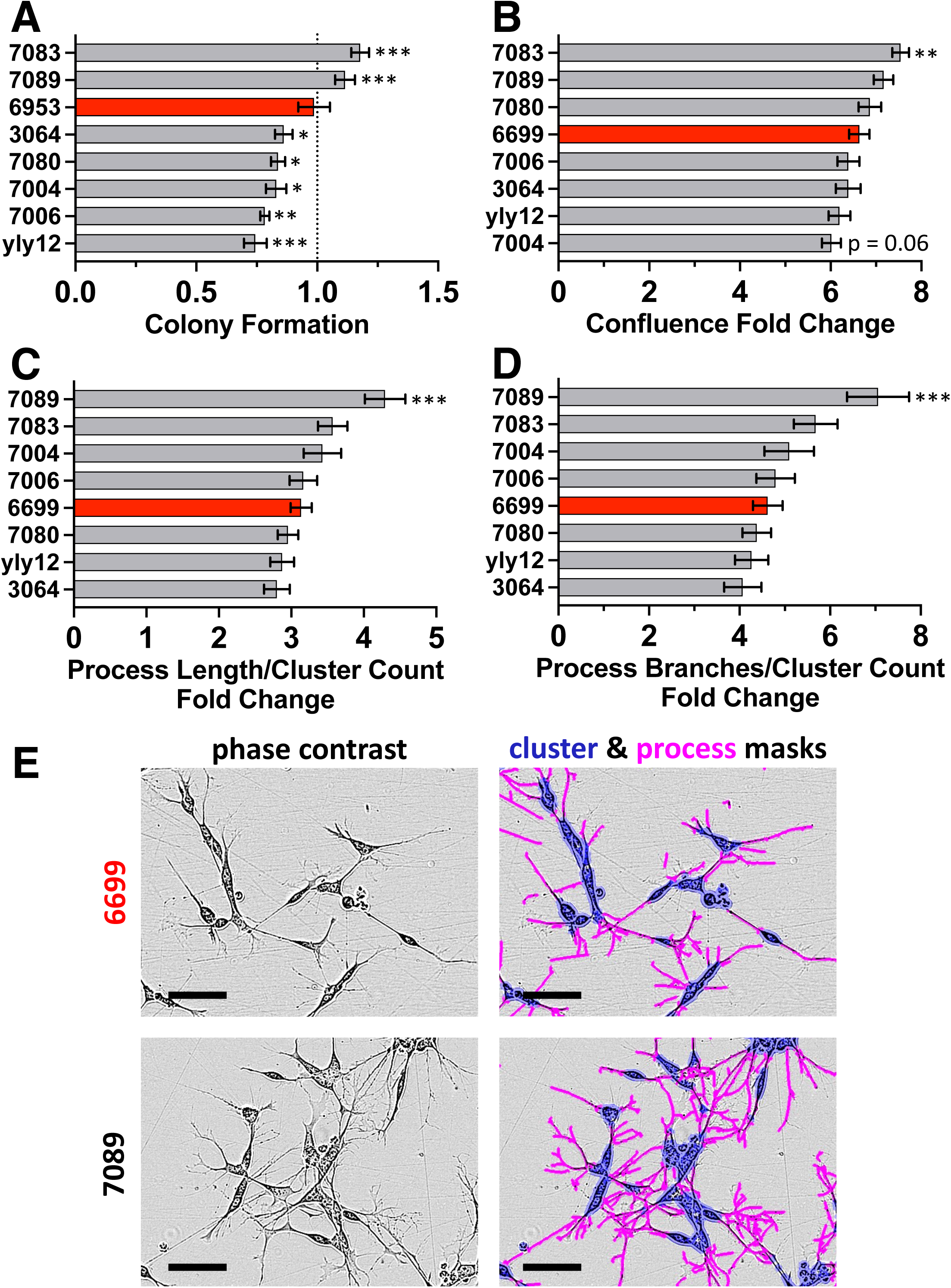
Aptamer treatment effects in cell culture. A) Daily treatment with 200 nM aptamers (gray) identifies those that promote or inhibit SH-SY5Y colony formation when compared to identical treatment with negative control oligonucleotides (red). Colony formation is compared to treatment with control oligonucleotides. Statistical significance is shown for comparison with 6953 (*: p<0.05, **: p<0.01, ***: p<0.001) by one-way ANOVA. Data are represented as mean ± SEM for biological replicates (n=4). B-D) Daily treatment with 350 nM aptamers (gray) identifies aptamers that promote increased confluence (B), neurite length per cluster count (C), and neurite branches per cluster count (D) when compared to identical treatment with a negative control oligonucleotide (red). Statistical significance is shown for comparison with 6699 (*: p<0.05, **: p<0.01, ***: p<0.001) by one-way ANOVA. Data are represented as mean ± SEM for biological replicates (n=46-48). E) Example regions of images quantified for C and D. The cluster mask is shown in blue, and the process mask is shown in pink. Scale bars are 100 µm.

The colony formation assay measured response to aptamer treatment on sparsely plated model cells over an a timeframe of 14 days. We tested aptamer effects on more densely plated cells over a shorter period of time (5 days) to prevent the culture reaching maximum confluence. Cell confluence was measured after 5 days of treatment, and while the general trend was the same as seen in the colony formation assay, only 7083 showed a statistically significant increase in confluence compared to a negative control oligonucleotide (Fig. 5B). The observed reduction in growth over a shorter treatment window reflects the time required to induce differentiation, typically requiring weeks rather than days.^33–35^ Since adherent SH-SY5Y cells in culture have short neurite-like processes, we quantified the effect of aptamer candidates on fold change in process length per cell cluster count, and process branches per cell cluster count. Aptamer 7089 showed a statistically significant increase in both morphological measurements when compared to negative control 6699. This is remarkable given the typical timeline of neural differentiation in cell culture (Fig. 5C-D). Phase contrast images and corresponding process and cluster masks are shown in Figure 5E.

The biological activity findings prompted followup with an RNA-seq study to determine gene expression changes in undifferentiated SH-SY5Y cells after 5 days of treatment with a subset of aptamers. All seven candidate aptamers that affected colony formation induced statistically significant changes in gene expression when compared with the negative control oligonucleotide. Aptamer 7004 showed the most pronounced response (Fig. 6A and Fig. S10). Gene Ontology (GO) analysis of up- and down-regulated genes in 7004-treated cells revealed many GO terms related to neuronal function (Fig. 6B-C). It is notable that many GO terms associated with metabolism and growth were significant when analyzing both the downregulated and upregulated genes from 7004-treated samples (Fig. S11 and Fig. S12). Gene Set Enrichment Analysis (GSEA) revealed two hallmarks from the 7004 treated sample data: oxidative phosphorylation and DNA repair (Fig. 6D and Fig. S13). Several GO terms also appear to be associated with oxidative phosphorylation when the genes upregulated by 7004 treatment were analyzed (Fig. 6E), while no GO terms were associated with DNA repair. Differential expression of neuronal genes resulted in significant GO pathways including synapse structure, axonogenesis, neurogenesis, and oxidative phosphorylation. Such changes have previously been associated with neuron differentiation.^34^ Notably, these differentiation-related changes were observed just 5 days after aptamer treatment, while negative control oligonucleotides treatment induces no such response. While the other six aptamers showed statistically significant changes in gene expression levels, GO analysis and GSEA did not reveal any significant terms. These other aptamers might be activating signaling networks that are not detected by gene expression alone at this particular treatment concentration and time point. These results constitute the first evidence a rapid biological response is elicited in a neuron model by sequence-specific aptamer interactions.

**Fig. 6.**
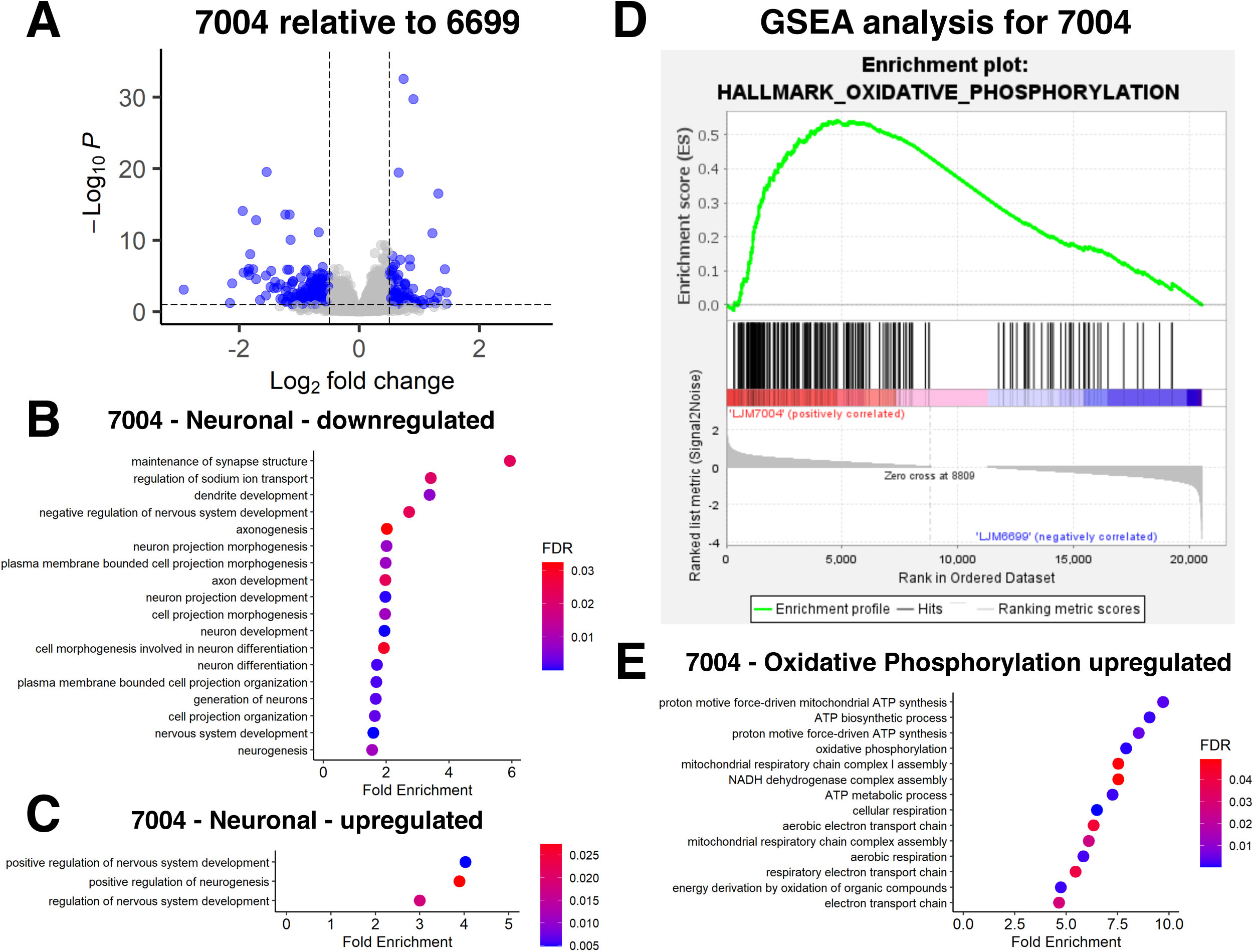
Gene expression changes after SH-SY5Y treatment with 7004. A) RNA-seq analysis of total RNA collected from cultures treated with aptamer (7004) or control oligonucleotide (6699). Log_2_ fold change was calculated as a ratio of 7004:6699. B-C) Neuronal GO terms resulting from analysis of downregulated (B) and upregulated (C) genes after treatment with 7004. D) Gene Set Enrichment Analysis (GSEA) revealed enrichment in oxidative phosphorylation for 7004. E) Gene Ontology (GO) terms associated with oxidative phosphorylation were upregulated after treatment with 7004.

## Discussion

This work describes a workflow for the selection and characterization of DNA aptamers with the potential to elicit biological responses in model neurons. This process led to the collection of 41 DNA aptamers displaying varying degrees of affinity for growing and differentiated SH-SY5Y cells. Of this collection, almost 20% (7/41) were found to induce changes in SH-SY5Y cell colony formation and gene expression detectable by RNA sequencing. While we have used a series of low-throughput biological assays in our workflow, we envision possible improvements. For example, a high throughput workflow could be applied to a large number of selected aptamers identified by deep sequencing without any need for cell affinity confirmation or characterization. In the future, it would be appropriate to expand this concept to explore libraries of selected DNA aptamers for biological activity in approaches similar to those taken with small molecules.^34^ Further, a high throughput biological activity assay would also allow for varying levels of streptavidin multimerization to more easily be tested. We note that in the present work we successfully leveraged cell replication rates and colony formation by proliferating model neurons as a powerful aspect of our screen for biological activity. This strategy is obviously limited to proliferating cells prior to differentiation.

Our results indicate that 7004 triggers biological activity that is in some ways reflective of neuron differentiation. Some responses to 7004 treatment are consistent with maturation of neuronal processes, most strongly supported by a shift towards oxidative phosphorylation. 7004 is therefore a candidate for further study as a neuroregenerative therapeutic agent potentially capable of promoting differentiation of induced pluripotent stem cells into mature neuronal cells.

Treatment of neurodegenerative diseases may require lifelong therapy and applications such as supporting nerve grafts after surgery also require extended periods of time. A hydrogel delivery system has been shown to provide a sustained release of aptamer.^36^ This hydrogel system suggests a slow-release approach to deliver aptamers to neural targets over weeks to months required for regeneration of injured nerves.

SH-SY5Y cells are often used as model neurons and therefore this collection of aptamers may find other useful applications in neuroscience. SH-SY5Y cells are also a useful model of neuroblastoma tumors, so this same collection of aptamers may possibly be useful leads for anti-proliferative therapeutics or cell-targeted drug delivery agents against neuroblastoma.

This work demonstrates a plausible workflow for future DNA aptamer selections against living cells or tissues to identify candidates that can then be screened for desired biological activity. We envision possible future studies in which aptamers are selected against various mature neurons to identify neuroregenerative oligonucleotides. Induced pluripotent stem cell-derived neuron models can be generated from patient-derived cells to identify aptamers that act specifically on diseased target cells. Further, aptamers capable of promoting specific differentiation outcomes might be identified by performing selections against naïve induced pluripotent stem cells.

## Materials and Methods

### Cell culture

SH-SY5Y cells were cultured in DMEM:F12 media (Gibco, 11330057) with 10% FBS (R&D Systems, S11550H) and 1% penicillin-streptomycin (Gibco, 15140122). HeLa and HEK293T cells were cultured in DMEM media (Gibco, 11965118) with 10% FBS (R&D Systems, S11550H) and 1% penicillin-streptomycin (Gibco, 15140122). These cells were maintained at 20% oxygen and 5% CO_2_.

### Buffers

Wash buffer is DPBS (Gibco, 14190-144) containing 5 mM MgCl_2_ and 4.5 g/L glucose. Binding buffer is containing 5 mM MgCl_2_, 4.5 g/L glucose, 100 µg/mL tRNA, and 100 µg/mL BSA. 2× PK buffer contains 100 mM Tris-HCl (pH 7.5), 200 mM NaCl, 2 mM EDTA, and 1% SDS. 0.5× TBE buffer contains 45 mM Tris-borate, pH 8.0 and 1 mM EDTA.

### Oligonucleotides

DNA library, primers, and aptamers were purchased from Integrated DNA Technologies (Coralville, IA) with standard desalting. All sequences and abbreviations are provided in Table S1.

### DNA aptamer selections

Three different selections (S1-S3) were performed. The protocol described below notes variations between selection conditions.

SH-SY5Y cells were cultured to ∼80% confluency on untreated 10-cm tissue culture dishes (Falcon, 353003) (S1: 4 million plated 24 h prior for first round and 2.25 million for subsequent rounds, S2 and S3: 2.5 million plated 48 h in advance with a media exchange at 24 h). Negative selection cells were cultured to ∼80% confluency as above (S1: no negative selection for rounds 1-2 and then plated 1 million NIH/3T3 cells 24 h prior for subsequent rounds, S2: no negative selection for rounds 1-2 and then plated 250,000 HeLa cells 48 h prior for subsequent rounds, S3: 1.5 million HEK293T cells for the first round and 750,000 cells for subsequent rounds plated 48 h in advance). 80-nucleotide (nt) naïve DNA libraries (40 central random nt flanked by blocks of 20 defined nts for PCR priming) were purchased from Integrated DNA Technologies (Coralville, IA) with standard desalting. For round 1, 1 nmol (∼6×10^14^ molecules) random DNA library (S1 and S2: library 6014, S3: library 6857) was prepared in 2 mL wash buffer, heated at 90°C for 5 min, and snap cooled on ice. Non-specific competitors were then added [S1: fragmented salmon sperm DNA (100 µg/mL) and BSA (1 µg/mL), S2 and S3: tRNA (100 µg/mL) and BSA (100 µg/mL)]. Subsequent rounds employed DNA library recovered and purified from large-scale (2 mL) PCR amplification (S1: 200 pmol for rounds 2-11 and 100 pmol for rounds 12-17, S2 and S3: 100 pmol). Media was aspirated from negative selection cells (when applicable) and the dish was rinsed once with DPBS. The library solution was added to the negative selection cells and incubated (S1: 4°C for 30 min in rounds 3-4 and for 60 min in rounds 5-17; S2: 37°C for 30 min in rounds 3-17; S3: 37°C for 30 min). Media was aspirated from SH-SY5Y cells and the dish was rinsed once with DPBS. When indicated, library solution was transferred from the negative selection cells to the SH-SY5Y cells and incubated (S1: 4°C for 60 min, S2: 37°C for 30 min, S3: 37°C for 30 min). For selection 1, SH-SY5Y cells were scraped into binding buffer, collected by centrifugation, and washed three times by centrifugation (500×g for 5 min). For selections 2 and 3, SH-SY5Y cells were washed on the dish three times with wash buffer, scraped into wash buffer, and collected by centrifugation (500×g for 5 min). The supernatant was removed, and the cell pellet was resuspended in 500 µL wash buffer. Cells were lysed by heating at 95°C for 10 min. S1 and S2: debris was removed by centrifugation at 13,100×g for 5 min, and the supernatant containing the recovered library was moved to a fresh tube. S3: the lysate was extracted with an equal volume of phenol:chloroform:iso-amyl alcohol (25:24:1) (VWR, 97064-692), and the upper aqueous phase was then precipitated from ethanol.

For the first selection round the cell pellet was lysed in water rather than wash buffer and the extract from the entire cell pellet was used as the template for 1 mL of PCR with the following reagent volumes: 500 µL round 1 lysate, 100 µL 10× PCR buffer, 80 µL 2.5 mM dNTPs, 100 µL 1 mg/mL BSA, 80 µL 50 mM MgCl_2_, 50 µL forward primer (S1 and S2: 5304, S3: 6638) (10 µM stock), 50 µL reverse primer (S1 and S2: 6015, S3: 6858) (10 µM stock), 20 µL water, 20 µL Taq (Invitrogen, 10342178). The mixture was split into 100 µL aliquots and subjected to thermal cycling [S1 and S2: 95°C, 60s; 8×(95°C, 30s; 70°C, 90s), S3: 94°C, 60s; 8×(94°C, 30s; 52°C, 35s; 72°C, 30s)].

The lysate (or the PCR product from the first round) was used directly as a template for analytical PCR to determine the optimum number of cycles to be used in a large-scale PCR. The following reagent volumes were used: 200 µL template, 200 µL 10× PCR buffer, 160 µL 2.5 mM dNTPs, 200 µL 1 mg/mL BSA, 160 µL 50 mM MgCl_2_, 200 µL forward primer (S1 and S2: 5304, S3: 6638) (5 µM stock), 200 µL reverse primer (S1 and S2: 6015, S3: 6858) (5 µM stock), 652 µL water, 28 µL Taq. The mixture was split into 100 µL aliquots and subjected to thermal cycling following the protocol described above with the optimum number of cycles as determined with analytical PCR followed by sample electrophoresis.

Following amplification, aliquots were recombined. 3M NaOAc was added and mixed (0.1 volumes). Nucleic acids were precipitated by the addition of 2.5 volumes of ethanol, mixing, incubation on dry ice for 15 min, and centrifugation at 17,000×g for 15 min. The pellet was washes with 70% ethanol followed by centrifugation at 17,000×g for 5 min. The pellet was air dried, resuspended in 40 µL water, combined with 160 µL deionized formamide, heated at 90°C for 5 min, and loaded onto a 10% denaturing polyacrylamide gel (7.5 M urea, 19:1 acrylamide:bisacrylamide) and subjected to electrophoresis for 2.5 h at 600 V (26.25 V/cm).

Bands were visualized with a handheld UV lamp. The higher mobility DNA band containing the fluorescent single stranded library was excised using a clean razor blade. This material was cleaved into small cubes and eluted overnight at 37°C in 500 µL 2× PK buffer on an end-over-end rotator. The supernatant was extracted against an equal volume of phenol:chloroform:iso-amyl alcohol (25:24:1) (VWR, 97064-692). The upper aqueous phase was then precipitated from ethanol as described above. The pellet was resuspended in 100 µL water. The concentration of the library was estimated using a molar extinction coefficient at 260 nm (S1 and S2: 881,375 M^-1^cm^-1^, S3: 769,475 M^-1^cm^-1^).

### Monitoring library recovery

qPCR reactions (30 µL) were performed with the following reagent volumes: 15 µL PerfeCTa SYBR Green FastMix for IQ (Quantabio, 95071-012), 3 µL 5 µM forward primer, 3 µL 5 µM reverse primer, 7.5 µL water, 1.5 µL template. Samples were transferred to a CFX96 Touch Deep Well Real-Time PCR Detection System (Bio-Rad Laboratories, 1854095) and subjected to thermal cycling using a protocol of 95°C, 30s; 40×(95°C, 15s; 51°C, 30s; 72°C, 30s), and interrogating SYBR Green fluorescence after the anneal step of each cycle as previously described.^26^ Raw fluorescence data were then analyzed using CFX Maestro™ software.

### Aptamer library sequencing and analysis

DNA libraries recovered after each selection round described above and the original, naïve library were subjected to PCR with the same protocol described above but with unmodified primers (S1 and S2: 5505 and 6160, S3: 6708 and 6895). The previously determined optimum number of cycles was used for each library to obtain unmodified, duplex DNA. PCR product size and quality were assessed by sample analysis by electrophoresis through 10% polyacrylamide gels with post-staining using SYBR Gold (Invitrogen, S11494) at this point and just before sequencing. MinElute spin columns (Qiagen, 28204) were used to purify the PCR products. A Qubit HS Duplex DNA Quantification kit (Invitrogen, Q32851) and Qubit 3.0 Fluorometer (Invitrogen, Q33216) were used to quantify the concentration of the purified PCR product. For each library, 10 ng was used as input into the NEBNext Ultra II DNA Library Prep with Sample Purification Beads (NEB, E7103S). Volumes recommended by the manufacturer were halved, and in the purification of adaptor-ligated DNA (step 3), Qiagen MinElute columns were used in place of beads. Libraries were barcoded using NEBNext Multiplex Oligos for Illumina (Index Primers Set 1 and Set 2) (NEB, E7335S and E7500S). Paired end sequencing was performed (S1 and S2: 150 cycles, S3: 100 cycles) on a single lane of an Illumina MiSeq instrument with 30% PhiX spike.

Usearch was used to merge paired end reads. Reads with a total quality score >0.5 were discarded.^37^ SeqKit was used to create uniform forward reads.^38^ AptaSUITE was used to filter any reads that did not contain both the forward and reverse primers within an error of 3 bases. AptaSUITE was also used to rank aptamers by their abundance and cluster the aptamers by sequence similarity (AptaCluster).^27^

### Immunofluorescence staining and IncuCyte SX5 imaging

Cells were washed with wash buffer. Aptamer solution (200 nM aptamer in wash buffer) was heated at 95°C for 5 min and snap cooled on ice. Final aptamer solution (200 nM aptamer in wash buffer) was added to cells for 1 h at 37°C. Cells were washed three times with wash buffer and incubated with 3.7% formaldehyde for 15 min at room temperature. Cells were washed twice with wash buffer and incubated with Streptavidin, Alexa Fluor 647 conjugate (Invitrogen, S21374) secondary stain at 1:500 dilution in DPBS for 1 h at room temperature. Cells were washed three times with DPBS and imaged with IncuCyte SX5 (Sartorius) at 20× magnification. Quantification of aptamer staining was analyzed using IncuCyte Basic Analyzer software. Confluence was analyzed under the Classic Confluence segmentation, adjustment set to 0.1 background-to-cell ratio with cleanup settings for ‘Hole Fill’ at 50.00 µm^2^ and ‘Adjust Size’ at 3 pixels. Aptamer staining was quantified using the Surface Fit segmentation routine with a threshold set to 1.5000 NIRCU. Individual images were then filtered for confluence levels (undifferentiated SH-SY5Y: minimum of 6.5%; differentiated SH-SY5Y: minimum of 13% and maximum of 85%). Blinded image data were also filtered for excess stain artifacts that impacted the fluorescence intensity measurement (undifferentiated SH-SY5Y: n=34-78 replicate image fields per condition in multiple replicate wells; differentiated SH-SY5Y: n=32-76 replicate image fields per condition in multiple replicate wells). The metric shown in figures as ‘Relative Fluorescence’ was calculated as the integrated fluorescent intensity per image relative to the percent confluence.

### SH-SY5Y Differentiation

Differentiation of SH-SY5Y cells used an optimized high-throughput protocol.^32^ Prior to plating, cells were thawed and kept in a basic growth medium: DMEM:F12 (Gibco, 11330057) with 10% FBS (R&D Systems, S11550H), 1% penicillin-streptomycin (Gibco 15140122) and glutamax-I (Gibco, 35050061). Cells were seeded in the basic growth medium on dishes coated with 10 µg/mL Laminin (Corning, 354232). The next day (Day 1 of differentiation) the medium was changed to Stage I media: DMEM with 2.5% FBS, 1% penicillin-streptomycin, glutamax-I, and 10 µM retinoic acid (Millipore Sigma, 554720). On day 4, medium was again changed with Stage I media. On days 6 and 9, the medium was changed to Stage II media: Neurobasal-A media (Gibco, 10888022) with 1% penicillin-streptomycin, glutamax-I, 20 mM KCl (Sigma, S9638), 50 ng/mL brain-derived neurotrophic factor (R&D Systems, 11166BD010), and B-27 (ThermoFisher Scientific, A3582801). Cells were used after day 11 of the differentiation process.

### Colony formation assay

SH-SH5Y cells were plated at 750 cells per well in 24-well plates (Falcon, 353047) and allowed to adhere for 24 h. Aptamer stock solutions for cell treatment (10× final indicated concentration) were prepared by heating and snap cooling in wash buffer and stored at -20 °C or 4 °C throughout the duration of the studies. Stock solutions (40 μL) were combined with growth media (360 μL) to reach the indicated final aptamer concentration for daily treatments. Growth media was replaced daily with 400 μL fresh media containing aptamer. Treatment continued for 13-14 days.

Media was then aspirated from cells followed by gentle washing with PBS. Cells were then fixed in 100% methanol for 10 min at -20 °C and immediately stained in 0.5% crystal violet (Sigma, C6158) for 10 min at -20 °C. Cells were washed 3-5 times with water to remove excess staining solution before quantification of colony formation.

### Optimizing biotinylated aptamer-streptavidin multimer stoichiometry

Stoichiometries of multimers of biotinylated aptamers bound to streptavidin were determined by holding aptamer concentration consistent at 1 µM and adding varying dilutions of Streptavidin, Alexa Fluor 647 conjugate (Invitrogen, S21374) in wash buffer and incubating at room temperature for 1 h with end-to-end rotation. Multimer mixtures were analyzed by electrophoresis through 8% polyacrylamide gels in 0.5× TBE running buffer for 3.5 h at 250 V (26.25 V/cm) followed by imaging with an Amersham Typhoon laser-scanning platform (Cytiva).

### Confocal imaging

Undifferentiated SH-SH5Y cells were plated on 35 mm glass-bottom dishes (MatTek, P35G-1.5-14-C) at 50-70% confluency and incubated overnight at 37 °C. Differentiated SH-SH5Y cells were plated on the same dishes and differentiated as described above before staining.

Biotinylated aptamers were incubated with fluorescently labeled streptavidin as described above prior to staining in all high-resolution images demonstrating aptamer binding (2% dilution of streptavidin). Cells on glass-bottom dishes were then gently washed twice in wash buffer before incubation with streptavidin-conjugated aptamer solutions for 1 h at 37 °C. Cells were washed 3 times in wash buffer and then fixed in formaldehyde as performed in IncuCyte SX5 imaging studies.

Fixed cells were washed twice in PBS to remove excess formaldehyde and stained with Actin555 ReadyProbe (Invitrogen, R37112) diluted in PBS per manufacturer’s recommendation for 30 min at room temperature. Cells were then washed twice more in PBS and stained with 1 μg/mL DAPI (Roche, 10236276001) in PBS for 5 min at room temperature. Plates were washed 3 times with PBS before imaging. Images were collected on a Zeiss LSM 780 confocal microscope.

### Confluence and neurite measurements of aptamer-treated SH-SY5Y cells

SH-SY5Y cells were plated (50,000 cells/well) on 24-well plates (Falcon, 353047) and allowed to adhere for 24 h at 37°C. Aptamer stocks (10× concentration) were prepared in advance in wash buffer. Solutions were heated at 90°C for 5 min and snap cooled and stored at 4°C for the treatment period. Daily media changes took place by replacing half of the volume of growth media and dosing with aptamer to yield a final concentration of 350 nM in each well. This experiment continued for 5 media changes and cells were imaged every 12 h with the Incucyte SX5 instrument.

Images were analyzed using the Incucyte Neurotrack software module (Sartorius, 9600-0010) for neurite measurements and the Basic Analyzer for confluence measurements. In the Neurotrack analyzer, the ‘Cell-Body Cluster Segmentation’ mode was set to ‘Brightness’ with the ‘Segmentation Adjustment’ value set to 1. The Cleanup Parameters were set to zero, with the ‘Minimum Cell Width (µm)’ set to 7.00. Neurite Parameter filtering was set to ‘Best’, ‘Neurite Sensitivity’ at 0.5, and ‘Neurite Width’ at 1 µm.

Confluence was measured using the Basic Analyzer under the Classic Confluence segmentation mode with the Segmentation Adjustment set to 0.1. The cleanup settings were set to zero, ‘Hole Fill’ set at 50.00 µm^2^ and ‘Adjust Size’ set at 3 pixels.

### RNA Sequencing of Aptamer-Treated Cells

SH-SY5Y cells were plated (100,000 cells/well) on 6-well plates (ThermoFisher Scientific, 353046) and allowed to adhere for 24 h. Aptamer stock solutions were prepared at 3.5 µM by heating and snap cooling in wash buffer and stored at 4°C throughout the duration of the treatment. Daily treatments were done by exchanging ¾ of well volume with aptamer diluted to 350 nM in growth media. Treatments were carried out for 5 d before RNA extraction.

Growth medium was then removed and the PureLink RNA Mini Kit (ThermoFisher Scientific, 12183018A) was used to extract RNA according to manufacturer recommendations for adherent mammalian cells. Genomic DNA was fragmented by passing lysate through a 20-gauge needle attached to an RNase-free syringe 10 times. Samples were then subjected to an on-column PureLink DNase treatment (ThermoFisher Scientific, 12185010) to minimize genomic DNA contamination. Yield and sample quality were analyzed by nanodrop and Qubit RNA High Sensitivity Assay Kit (ThermoFisher Scientific, Q32852). Sample yields were ≥2 µg at ≥50 ng/µL with A260/A280 purity ratio between 1.8 and 2.2 as recommended by Azenta Life Sciences.

Isolated RNA samples were stored at -80°C and transferred into RNA Stabilization Tubes (Azenta Life Sciences, GTR5025-GW) prior to sample submission. 40 µL of isolated RNA was added to tubes containing stabilization matrix, mixed thoroughly and air dried in a biosafety hood for 24 h.

Isolated RNA was sequenced by Azenta Life Sciences. 50 million reads per sample were obtained in biological triplicates. Transcript sequences were aligned versus human genome assembly GRCh38.p14 and quantified with Salmon.^39,40^ Differential expression analysis was performed with DESeq2 with Benjamini and Hochberg false discovery rate correction.^41^ Volcano plots were generated with EnhancedVolcano.^42^

Gene Ontology (GO) analysis of up- and downregulated transcript sets was performed by the Gene Ontology Resource in release 2023-11-15.^43–45^ Gene Set Enrichment Analysis (GSEA) performed in GSEA_4.3.2 using transcript quantification from Salmon.^46, 47^

## Data Availability

All data supporting the statements and conclusions made by the authors are included in the figures and in the supplemental information.

## Acknowledgments

This work was supported by NIH grant R35GM143949 (LJM), DOD grant W81XWH-22-1-0313 (LJM), an NSF graduate fellowship (BW), and by the Mayo Clinic Graduate School of Biomedical Sciences. The authors thank members of the Maher and Staff laboratories. We acknowledge the Mayo Clinic Genome Analysis Core for their contributions.

## Author Contributions

Conceptualization: K.P., A.E.W., and L.J.M. Funding acquisition: A.E.W., N.P.S., and L.J.M. Investigation: J.R., K.P., B.W. Data analysis: J.R., K.P., B.W., S.C.L.H., L.J.M. Visualization: J.R., K.P., B.W. Writing – original draft: J.R., K.P. Writing – review & editing: J.R., K.P., B.W., S.C.L.H., A.E.W., N.P.S., L.J.M. All authors read and approved the final manuscript.

