## supplemental materials for "DNA aptamers that modulate biological activity of model neurons"

**Supplemental Figures and Tables: DNA aptamers that modulate biological activity of neuron models**

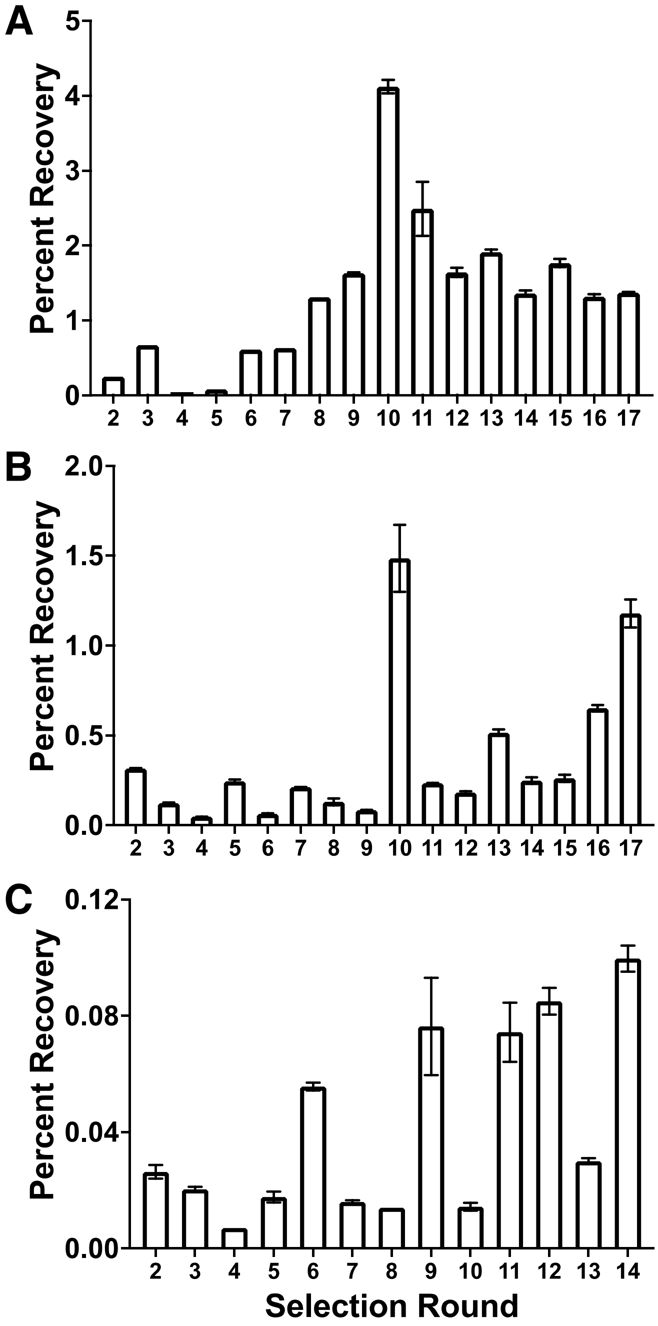


**Figure S1.** qPCR quantification of library recovery across selection rounds. A-C) Data from selections 1-3, respectively. Data are represented as mean ± SEM.

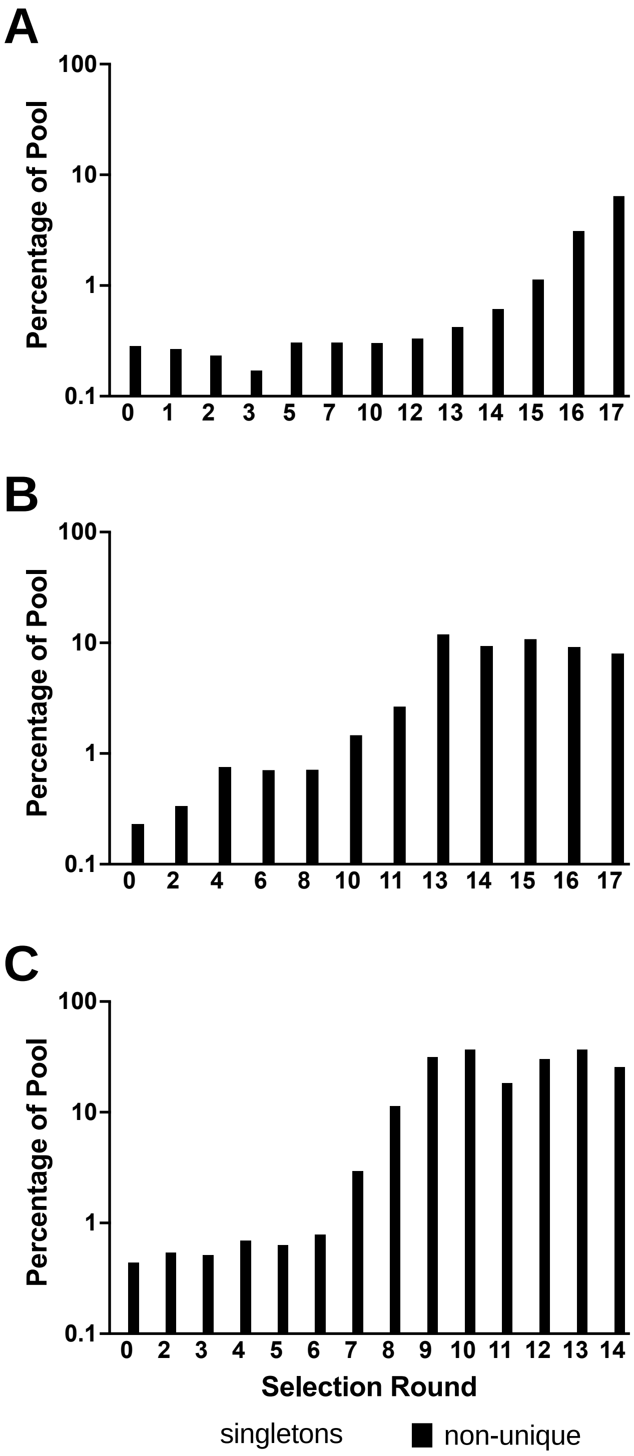


**Figure S2.** Library enrichment from deep sequencing data across selection rounds. A-C) Data from selections 1-3, respectively. “Singletons”: sequences found only once in deep sequencing data. “Non-unique”: sequences found more than once in deep sequencing data.

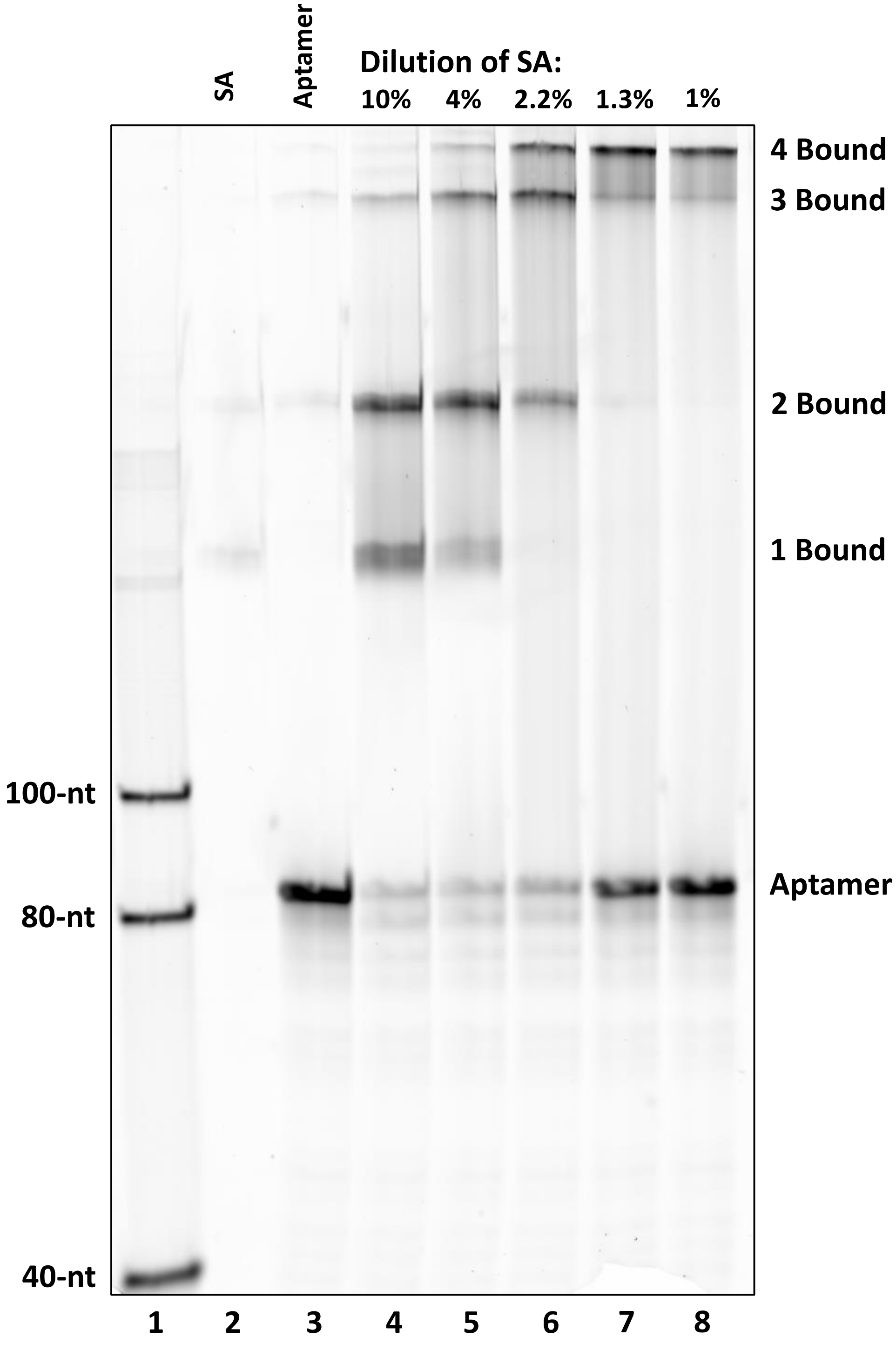


**Figure S3.** Optimizing biotinylated aptamer-streptavidin multimer conjugates. Native polyacrylamide gel shows fluorescein modified aptamer and aptamer-streptavidin complexes. Volume and aptamer concentration was held constant (1 μM), and Streptavidin Alexa Fluor 647 conjugate (Invitrogen, S21374) stock was added at varying dilutions, expressed as a percent dilution by volume. Lane 1: single-stranded DNA ladder. Lane 2: streptavidin only (SA). Lane 3: aptamer only. Lane 4-8: varying percent dilution by volume from stock Streptavidin, Alexa Fluor 647 conjugate.

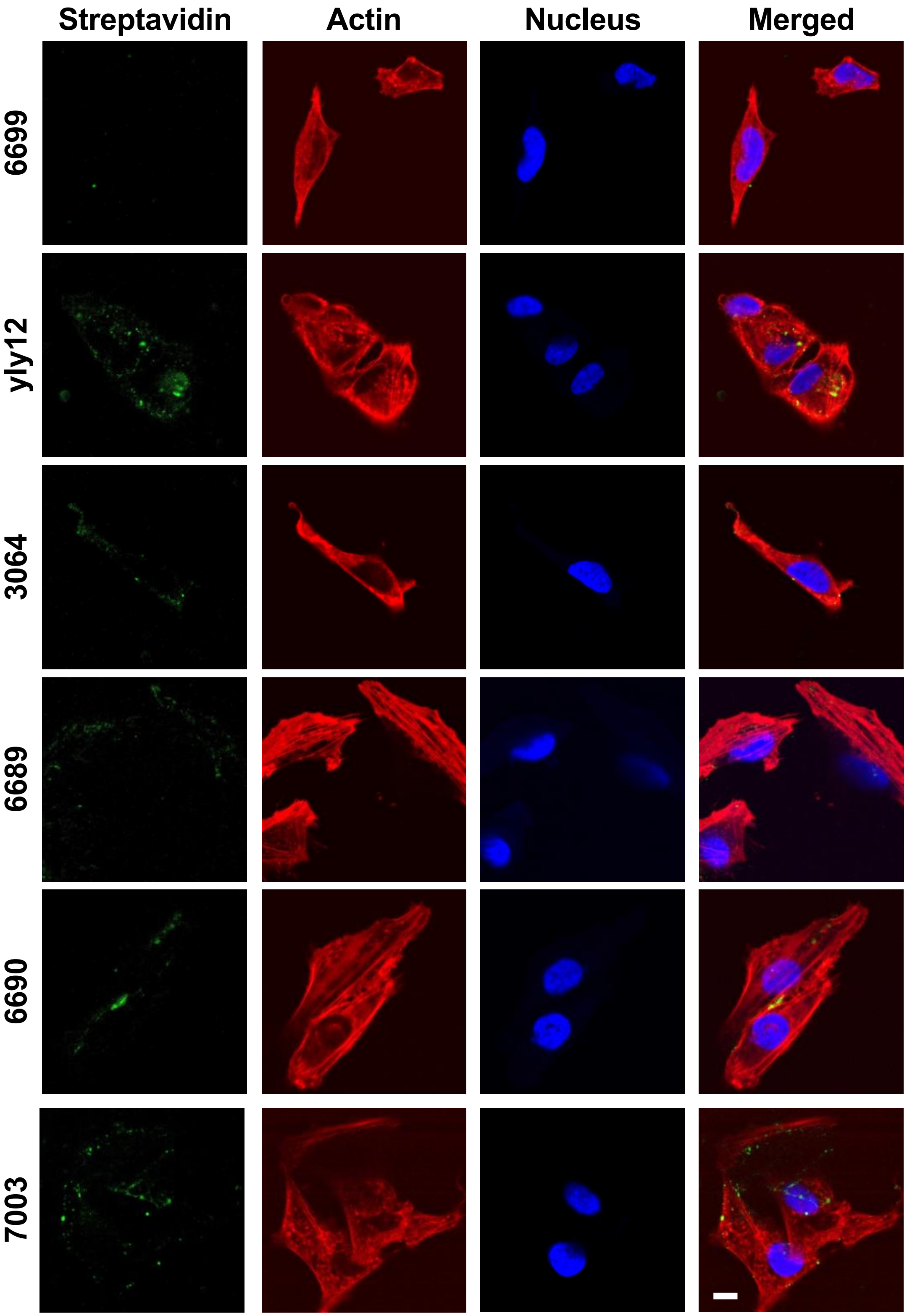


**Figure S4.** Fluorescence confocal imaging of aptamer staining on undifferentiated SH-SY5Y cells. Binding of biotinylated aptamers is detected by staining with Alexa Fluor 647-labeled streptavidin (green). Cells are detected by staining actin with fluorescently labeled phalloidin (red) and with DAPI (blue). 100× magnification, scale bar is 10 μm (shown in lower right image).

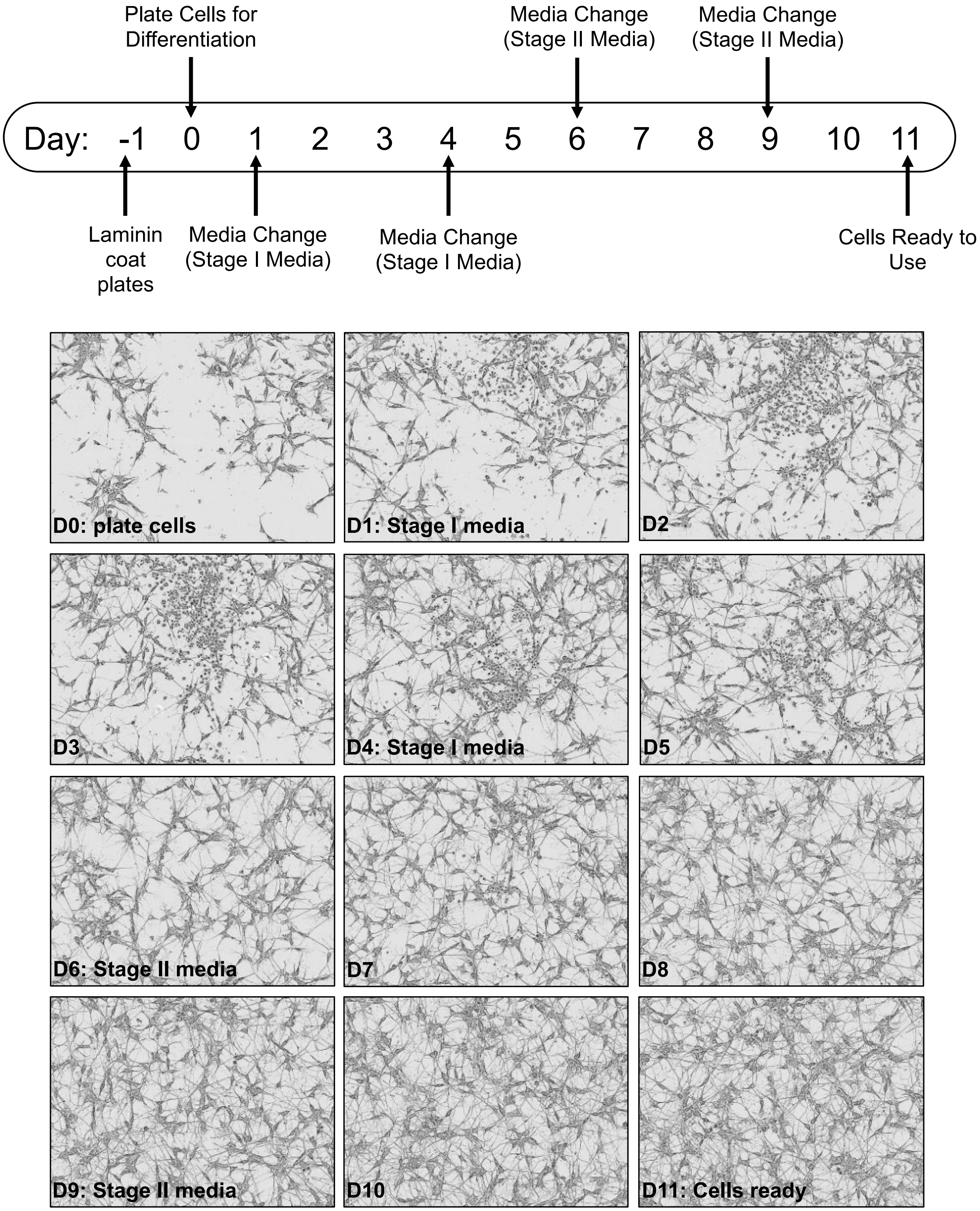


**Figure S5.** Differentiation of SH-SY5Y cells. SH-SY5Y differentiation is performed over 12 d with four media changes. Cells were incubated first in Stage I media, consisting of 10 µM retinoic acid, then in Stage II media, containing 50 ng/mL BDNF, to produce differentiated neuronal cells. Throughout the differentiation process, phase contrast images were obtained every 24 h. Cells continued to proliferate for a short time until fully differentiated, between day 7 and day 9. Images were acquired using Incucyte SX5 at 20× magnification.

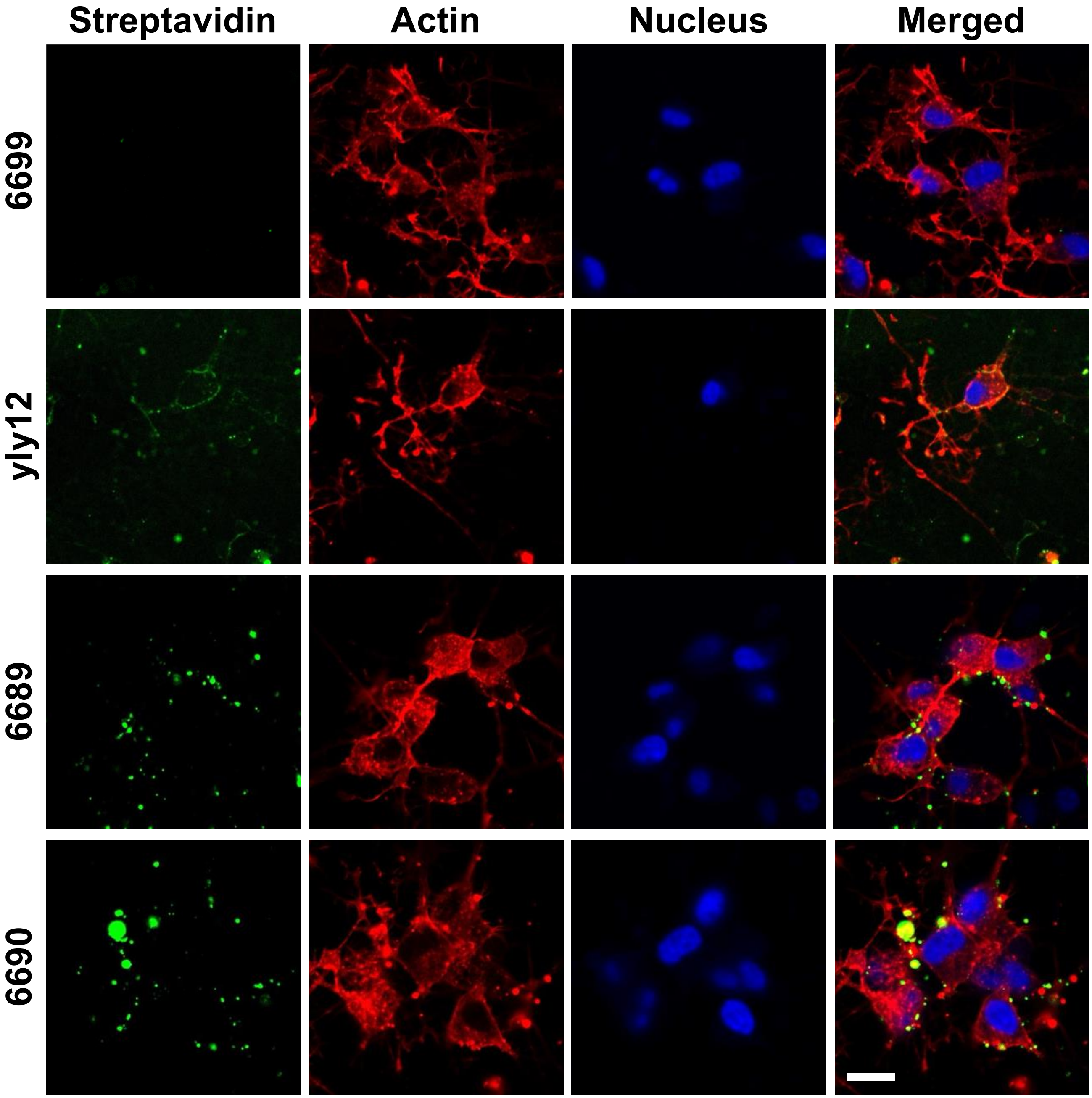


**Figure S6.** Fluorescence confocal imaging of aptamer staining on differentiated SH-SY5Y cells. Binding of biotinylated aptamers is detected by staining with AlexaFluor647-labeled streptavidin (green). Cells are detected by staining actin with fluorescently labeled phalloidin (red) and with DAPI (blue). 40× magnification, scale bar is 20 µm (shown in lower right image).

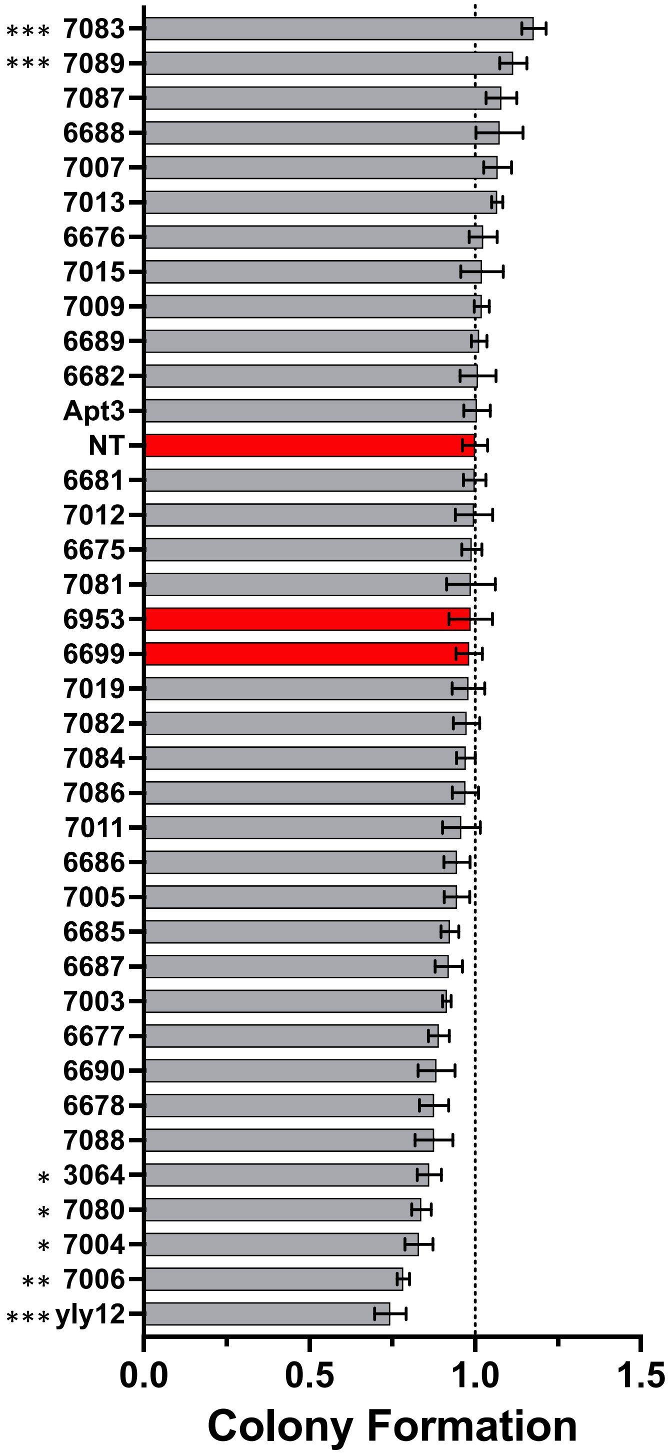


**Figure S7.** Aptamer treatment effects in colony formation assay. A) Daily treatment at 200 nM identifies aptamers (gray) that promote or inhibit SH-SY5Y colony formation when compared to identical treatment with negative control oligonucleotides (red). Colony formation efficiency is relative to NT (no treatment). Statistical significance is shown for comparison with control 6953 (*: $p<0.05$, **: $p<0.01$, ***: $p<0.001$) by one-way ANOVA. Data are represented as mean ± SEM for biological replicates (n=4).

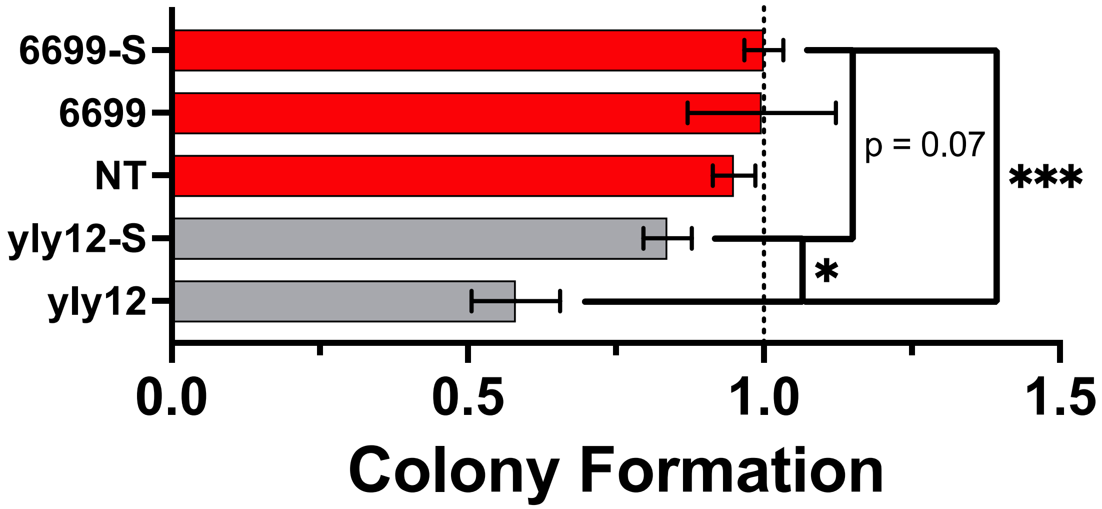


**Figure S8.** Multimerization of yly12 by streptavidin conjugation decreases its effect in colony formation assay. Daily treatment with 200 nM yly12 multimerized with streptavidin (yly12-S) did not statistically significantly inhibit SH-SY5Y colony formation when compared to identical treatment with negative control oligonucleotide 6699 with or without streptavidin. Colony formation efficiency is relative to NT (no treatment). Data are represented as mean ± SEM for biological replicates (n=4). Statistical significance was determined by one-way ANOVA (*: $p<0.05$, **: $p<0.01$, ***: $p<0.001$).

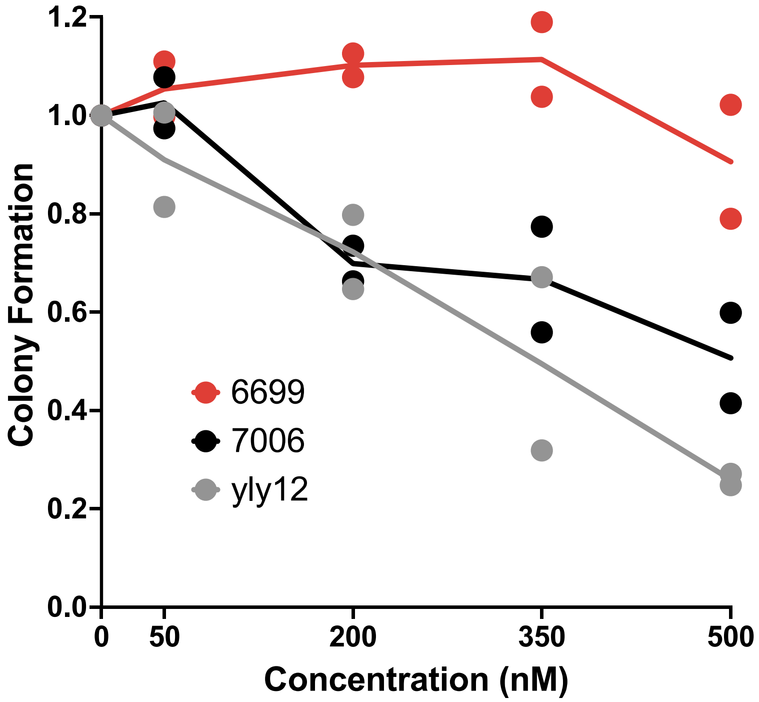


**Figure S9.** Aptamer dose-response tested in colony formation assay. Daily treatment with 50, 200, 350, or 500 nM aptamers [7006 (black) and yly12 (gray)] in culture media shows a dose-dependent inhibition of SH-SY5Y colony formation when compared to identical treatment with negative control oligonucleotide 6699 (red). Treatment with 6699 begins to show inhibition of colony formation at 500 nM. Colony formation efficiency is relative to NT (no treatment).

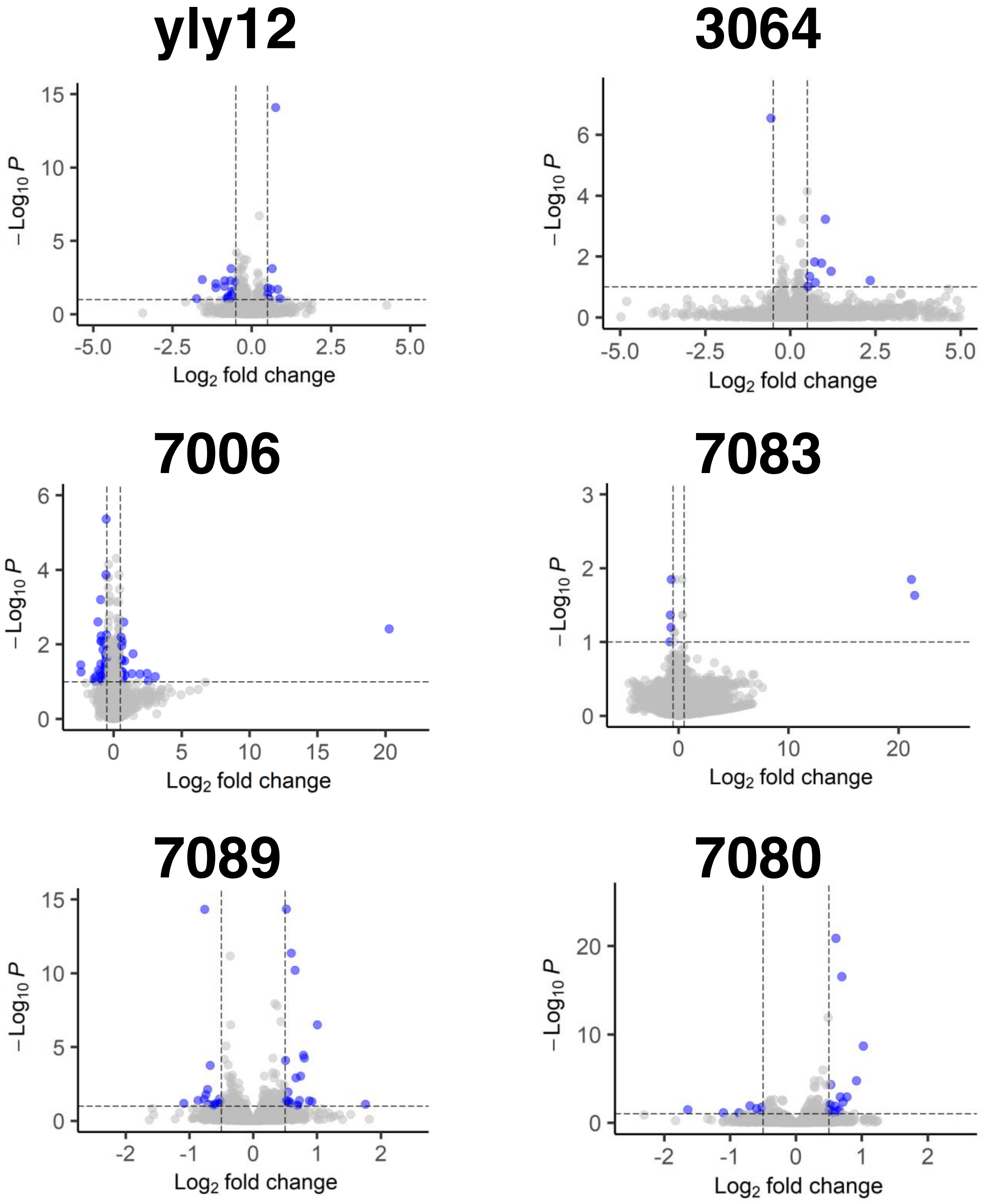


**Figure S10.** Gene expression changes after SH-SY5Y cells were treated with aptamers. RNA-seq analysis of total RNA collected from cultures treated with aptamer or control oligonucleotide (6699). $Log_2$(fold change) was calculated for the aptamer:control oligonucleotide ratio.

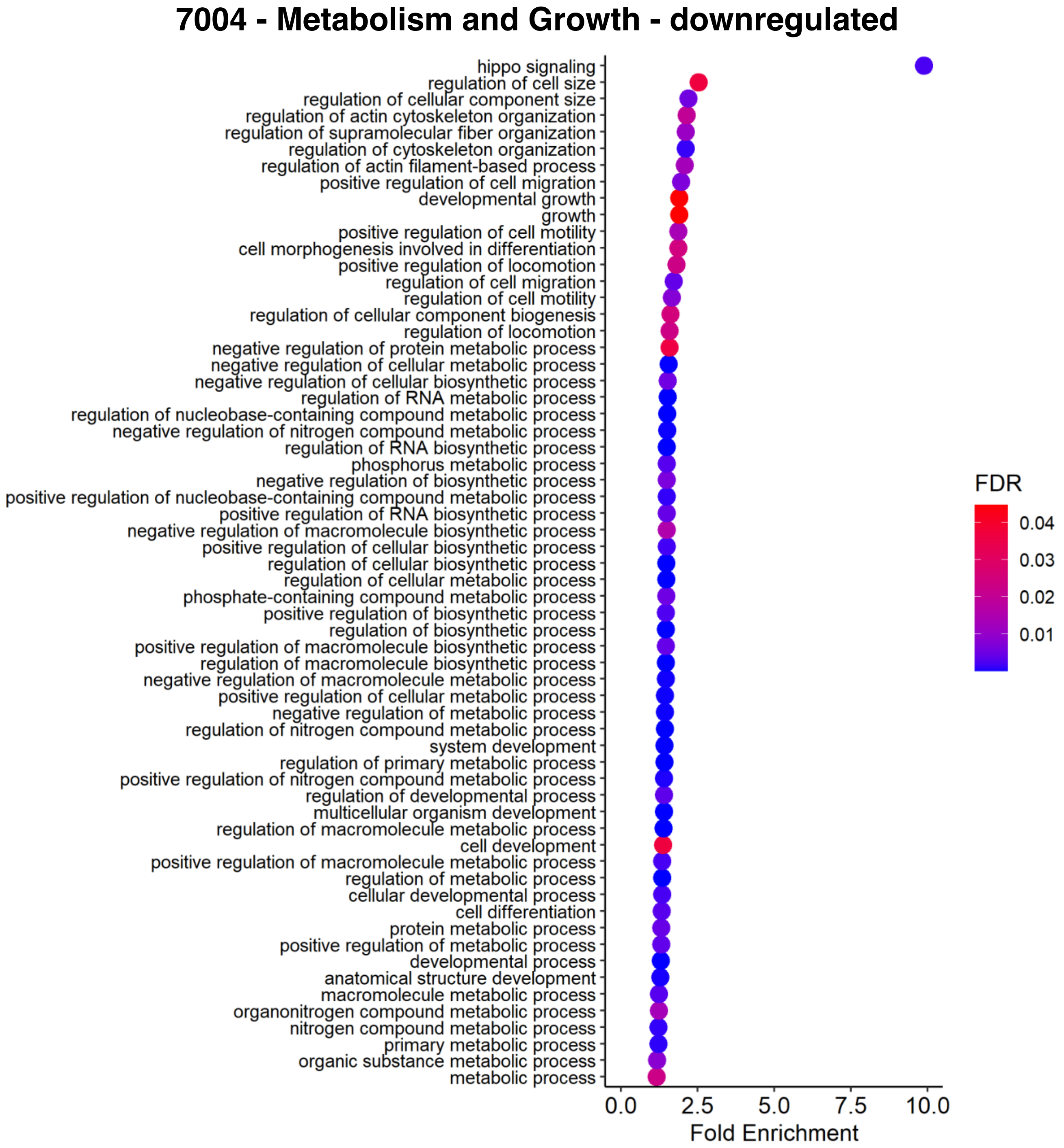


**Figure S11.** Metabolism- and growth-associated terms revealed in downregulated expression data after treatment with aptamer 7004. GO analysis reveals metabolism- and growth-associated terms when analyzing the downregulated genes after treatment with 7004.

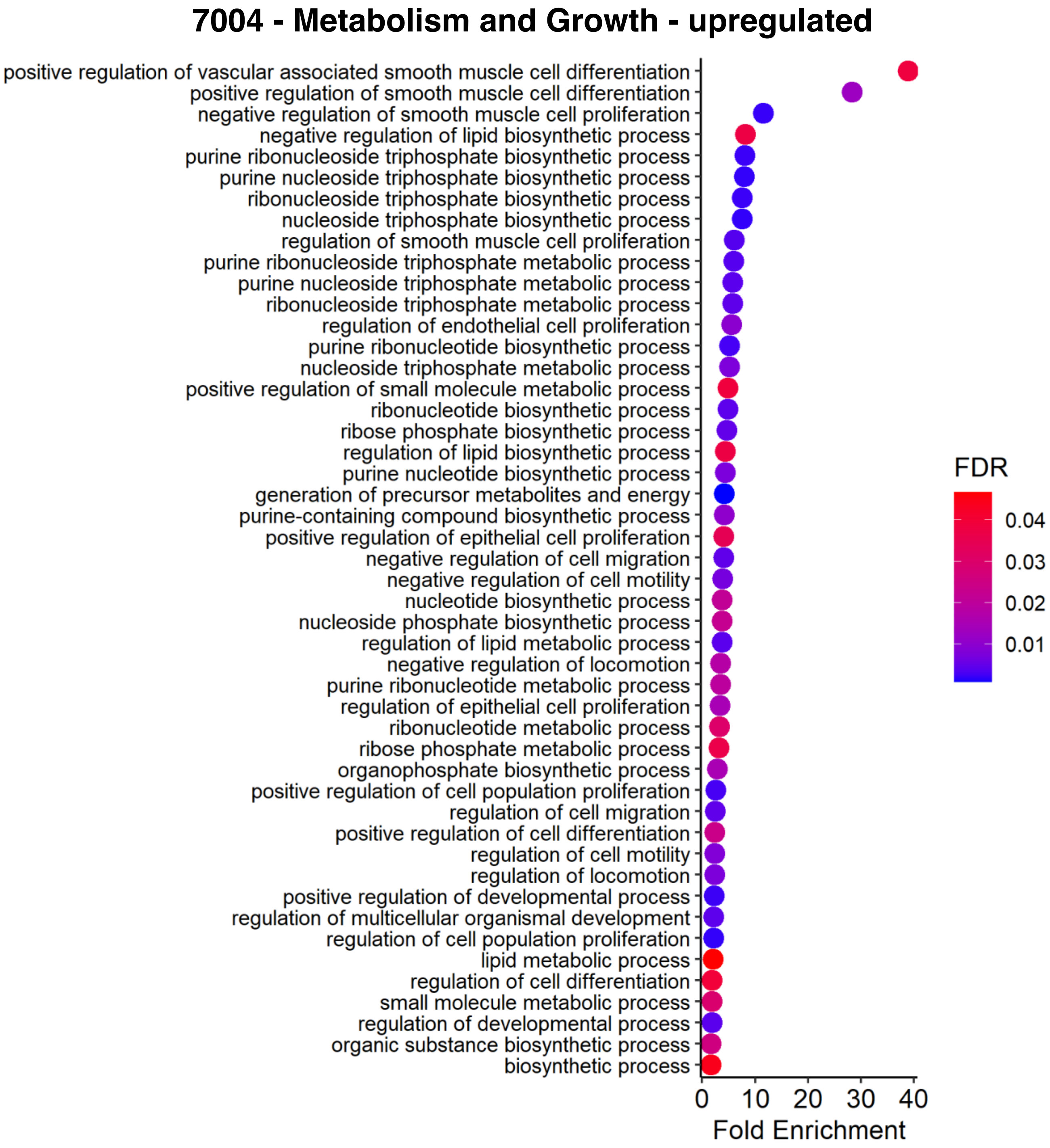


**Figure S12.** Metabolism- and growth-associated terms revealed in upregulated expression data after treatment with aptamer 7004. GO analysis reveals metabolism- and growth-associated terms when analyzing the upregulated genes after treatment with 7004.

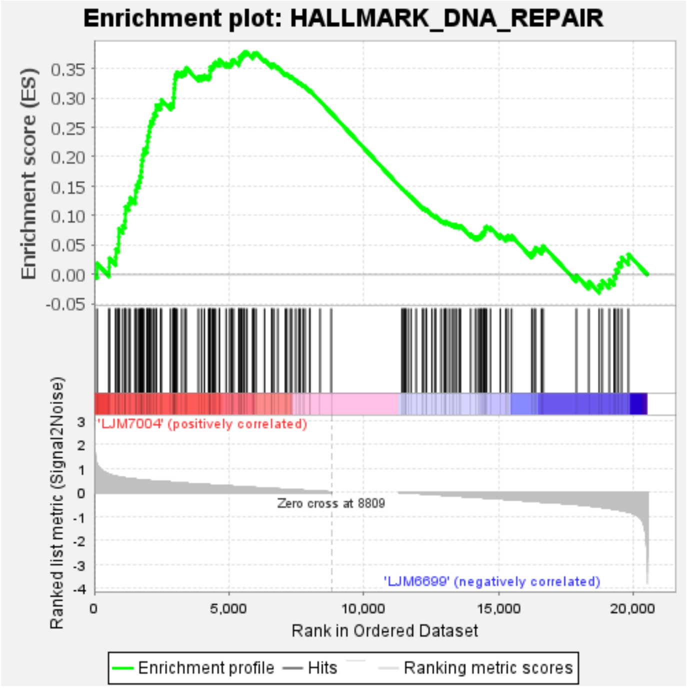


**Figure S13.** GSEA reveals enrichment in DNA repair after treatment with aptamer 7004.

**Table S1. Designation and primary structure of primers, libraries, and aptamers.**

| Serial number[1] | Oligonucleotide sequence[2] |
|---|---|
| 3064 | /56-FAM/GGGTCGGCGGGTGGGGTGGGAGGTGGTCTTGTCTCTGGGT/3BioTEG/ |
| 5304 | /56-FAM/+AG+AC+CA+GACCAGCTGATACCAGTCGTG |
| 5505 | AGACCAGACCAGCTGATACCAGTCGTG |
| 6014 | AGACCAGACCAGCTGATACCAGTCGTGN$_{40}$CCACGTAGCTCTCCAGCTCCACCTGA |
| 6015 | A$_{20}$/iSp9//iSp9/TCAGGTGGAGCTGGAGAGCTACGTGG |
| 6160 | TCAGGTGGAGCTGGAGAGCTACGTGG |
| 6202 | /56-FAM/AGACCAGACCAGCTGATACCAGTCGTGCCTCATAGACGATCACCGTTTCACC CATACGAAAGGTATACCACGTAGCTCTCCAGCTCCACCTGA/3BioTEG/ |
| yly12 (6214) | /56-FAM/AGGATAGGGGGTAGCTCGGTCGTGTTTTTGGGTTGTTTGGTGGGTCTTCTG/ 3BioTEG/ |
| 6638 | /56-FAM/TGCGTATTGACACATGCGTG |

6673 /56-FAM/AGACCAGACCAGCTGATACCAGTCGTGGTGCTATTTATAGAGTCTCAAGGGA
ACCGGACTTGAAACGCCACGTAGCTCTCCAGCTCCACCTGA/3BioTEG/

6674 /56-FAM/AGACCAGACCAGCTGATACCAGTCGTGTAGACAGAAGTGCTCTTGCCGACTC
TGGTATCGCATCTCACCACGTAGCTCTCCAGCTCCACCTGA/3BioTEG/

6675 /56-FAM/AGACCAGACCAGCTGATACCAGTCGTGGGCGCCCCGCGGTTAAAAGGAGATC
TATCCGGACCTATCACCACGTAGCTCTCCAGCTCCACCTGA/3BioTEG/

6676 /56-FAM/AGACCAGACCAGCTGATACCAGTCGTGGCAATGCTAGCATAGCGTCAATAAG
GTAACGGACGTAACACCACGTAGCTCTCCAGCTCCACCTGA/3BioTEG/

6677 /56-FAM/AGACCAGACCAGCTGATACCAGTCGTGGGCCGTTTAGGAGTTCATCAAGCTT
CAGCGGACTTGTAACCCACGTAGCTCTCCAGCTCCACCTGA/3BioTEG/

6678 /56-FAM/AGACCAGACCAGCTGATACCAGTCGTGGCGCGATGAAGCGCGTTCTAAGTGA
CAACGGACGGGCCACCCACGTAGCTCTCCAGCTCCACCTGA/3BioTEG/

6679 /56-FAM/AGACCAGACCAGCTGATACCAGTCGTGGATGGAAAAACCGGGTTGTGTAGAC
TGCAGAGACTGCCGACCACGTAGCTCTCCAGCTCCACCTGA/3BioTEG/

6680 /56-FAM/AGACCAGACCAGCTGATACCAGTCGTGGACCGTAGGGGCCAAGCAAGCTCAG
ACCGTAGGGACTGTGCCACGTAGCTCTCCAGCTCCACCTGA/3BioTEG/

6681 /56-FAM/AGACCAGACCAGCTGATACCAGTCGTGAAGAACCAGGGCGGAGGGCCTGTCT
CGATACGAACGTTAACCACGTAGCTCTCCAGCTCCACCTGA/3BioTEG/

6682 /56-FAM/AGACCAGACCAGCTGATACCAGTCGTGAAGTGAACCCGCGGACTGCAGAGAC
CACGGGAAGCCTCTCCCACGTAGCTCTCCAGCTCCACCTGA/3BioTEG/

6683 /56-FAM/AGACCAGACCAGCTGATACCAGTCGTGGTGGCAGACTGTAGAGACTGACACA
GGTGAATGACCGAATCCACGTAGCTCTCCAGCTCCACCTGA/3BioTEG/

6684 /56-FAM/AGACCAGACCAGCTGATACCAGTCGTGAAAAGAAATCTAGACGGCCGCGACT
AACTTGATAGAACGACCACGTAGCTCTCCAGCTCCACCTGA/3BioTEG/

6685 /56-FAM/AGACCAGACCAGCTGATACCAGTCGTGGAGCTGCATTCTATCCCTATCACAT
GTCACGACTGATCAACCACGTAGCTCTCCAGCTCCACCTGA/3BioTEG/

6686 /56-FAM/AGACCAGACCAGCTGATACCAGTCGTGTGGGAATCACGACTTCACCATCATT
TGTTTTTGCACGACTCCACGTAGCTCTCCAGCTCCACCTGA/3BioTEG/

6687 /56-FAM/AGACCAGACCAGCTGATACCAGTCGTGCGATAGCCCTTTAGGTGACTTTAAT
AACTGACTCCTTGGACCACGTAGCTCTCCAGCTCCACCTGA/3BioTEG/

6688 /56-FAM/AGACCAGACCAGCTGATACCAGTCGTGGTCCGAACGAGACATGCGCGCTAGC
CCTTTAGGACATTCTCCACGTAGCTCTCCAGCTCCACCTGA/3BioTEG/

6689 /56-FAM/AGACCAGACCAGCTGATACCAGTCGTGTGCACGACTGCCCCTCTAAAGCCTA
TGCACGACTGTCCGCCCACGTAGCTCTCCAGCTCCACCTGA/3BioTEG/

6690 /56-FAM/AGACCAGACCAGCTGATACCAGTCGTGATACCCCCTTGTGTGCGTGTTGTGT
GCCCATGTATTCTTTCCACGTAGCTCTCCAGCTCCACCTGA/3BioTEG/

6699 /56-FAM/AGACCAGACCAGCTGATACCAGTCGTGCATCAAGCCCCCGACCCGTAACGGA
AAGAATATGTATAACCCACGTAGCTCTCCAGCTCCACCTGA/3BioTEG/

6708 TGCGTATTGACACATGCGTG

6857 TGCGTATTGACACATGCGTG$N_{40}$TCGGTGAAGTAGTGTTACGC

6858 $A_{20}$/iSp9//iSp9/GCGTAACACTACTTCACCGA

6895 GCGTAACACTACTTCACCGA

6943 TGCGTATTGACACATGCGTGCGCTCGTCTTTGATGCGGGCTGTAGTGCGAAGTAGGTCTT
TCGGTGAAGTAGTGTTACGC/3BioTEG/

6944 TGCGTATTGACACATGCGTGTAGACAGGAGTGACGACCGTTCATATGTCCAATCTACTGA
TCGGTGAAGTAGTGTTACGC/3BioTEG/

| | |
|---|---|
| 6945 | TGCGTATTGACACATGCGTGCGAGCTAACCACGGACGATCTAGACTGTTGAGACTTGGCG TCGGTGAAGTAGTGTTACGC/3BioTEG/ |
| 6946 | TGCGTATTGACACATGCGTGGAGTTAGCGGACTGTAGAGACCACTAAATCGTTTAACCAC TCGGTGAAGTAGTGTTACGC/3BioTEG/ |
| 6947 | TGCGTATTGACACATGCGTGCAGACTGCAGAGACTGCCCATCCAATGCGCACAATACCAT CTCGGTGAAGTAGTGTTACGC/3BioTEG/ |
| 6948 | TGCGTATTGACACATGCGTGTAGGCCGGAGGGACTTCTAATGTTTGTCAATCTAAAGAGG TCGGTGAAGTAGTGTTACGC/3BioTEG/ |
| 6949 | TGCGTATTGACACATGCGTGCAGCCAATAGATAACACACAGAATCCGGACCGTAGGGCTC TCGGTGAAGTAGTGTTACGC/3BioTEG/ |
| 6950 | TGCGTATTGACACATGCGTGGTGTAGACGGAAGCGCGTTCTGCCCAAATTGTGGACCCCT TCGGTGAAGTAGTGTTACGC/3BioTEG/ |
| 6951 | TGCGTATTGACACATGCGTGCACATCACCGACCTACTACTTCTACCTTCCACACCACGCA TCGGTGAAGTAGTGTTACGC/3BioTEG/ |
| 6952 | TGCGTATTGACACATGCGTGCACCCCTCAAGCGTGGAGTTTTAAGGTTCACCCGGACTGT TCGGTGAAGTAGTGTTACGC/3BioTEG/ |
| 6953 | TGCGTATTGACACATGCGTGATTTAGAGTAAATCAACCTAGACTATCCGTGAGATACATC TCGGTGAAGTAGTGTTACGC/3BioTEG/ |
| 6954 | TGCGTATTGACACATGCGTGATCCTGCCCTTTTATAATAACTTCGCTCTAAGGATTTGCC TCGGTGAAGTAGTGTTACGC/3BioTEG/ |
| Apt3 (6965) | TAGGGAATTCGTCGACGGATCCTACCCGTTGCTGCAGGATCCTGAGATCGCCTCTGTCTG CAGGTCGACGCATGCGCCG/3BioTEG/ |
| 7003 | AGACCAGACCAGCTGATACCAGTCGTGGGATGCATCTCTCCTTGCATCGTTCGCCTCCAC CCGAACCCCACGTAGCTCTCCAGCTCCACCTGA/3BioTEG/ |
| 7004 | AGACCAGACCAGCTGATACCAGTCGTGGCACAAACGCCACTTTGCATCTAATCTCAACAT TCGATGGCCACGTAGCTCTCCAGCTCCACCTGA/3BioTEG/ |
| 7005 | AGACCAGACCAGCTGATACCAGTCGTGGGTCGTTCACCTCGCCGGTTTTAAGCCAACCCA GGCCATCCCACGTAGCTCTCCAGCTCCACCTGA/3BioTEG/ |
| 7006 | AGACCAGACCAGCTGATACCAGTCGTGATAGGGGCACGCCCACGGAAAAGTCAACCCGTC CACGGCACCACGTAGCTCTCCAGCTCCACCTGA/3BioTEG/ |
| 7007 | AGACCAGACCAGCTGATACCAGTCGTGCACATTCTCAACAATGCGGGTTCGGAACCGTCC CCGTTAACCACGTAGCTCTCCAGCTCCACCTGA/3BioTEG/ |
| 7008 | AGACCAGACCAGCTGATACCAGTCGTGTCAGCAATCCCTACTTTGCTCTGAACACACTGC GCTACACCCACGTAGCTCTCCAGCTCCACCTGA/3BioTEG/ |
| 7009 | AGACCAGACCAGCTGATACCAGTCGTGTGCATTTCTCCCAATGTGGTTATGGCCCCACGC TTTGCGCCCACGTAGCTCTCCAGCTCCACCTGA/3BioTEG/ |
| 7010 | AGACCAGACCAGCTGATACCAGTCGTGGTAATCCATCCCAAATGGATCCCGCTCACCAAC AGCGGCACCACGTAGCTCTCCAGCTCCACCTGA/3BioTEG/ |
| 7011 | AGACCAGACCAGCTGATACCAGTCGTGGAAAGCACGGACCCCAACTCCGGCACGCACCAC CGCTACCCCACGTAGCTCTCCAGCTCCACCTGA/3BioTEG/ |
| 7012 | AGACCAGACCAGCTGATACCAGTCGTGCGCACCCACTCGCGCAGGCGTAGACCCACCTCT CGCCCGACCACGTAGCTCTCCAGCTCCACCTGA/3BioTEG/ |
| 7013 | AGACCAGACCAGCTGATACCAGTCGTGTGACCAATTCCCTACTATTGCGTCACAAACTTA GCATACCCCACGTAGCTCTCCAGCTCCACCTGA/3BioTEG/ |
| 7014 | AGACCAGACCAGCTGATACCAGTCGTGACCGCTTTCACCAACAAACCGTCGACGTACTGA GCGTACCCCACGTAGCTCTCCAGCTCCACCTGA/3BioTEG/ |
| 7015 | AGACCAGACCAGCTGATACCAGTCGTGGTCTTACTGGAGGGCCATCCCCAATCATGGCCA GCGTACCCCACGTAGCTCTCCAGCTCCACCTGA/3BioTEG/ |

| | |
|---|---|
| 7080 | /56-FAM/CTGATACCAGTCGTGGGATGCATCTCTCCTTGCATCGTTCGCCTCCACCCGA |
| 7081 | /56-FAM/TCGTGGCACAAACGCCACTTTGCATCTAATCTCAACATTCGATGGCCACGTA |
| 7082 | /56-FAM/CCAGTCGTGGCAATGCTAGCATAGCGTCAATAAGGTAACGGACGTAACACCA |
| 7083 | /56-FAM/TCGTGAAGAACCAGGGCGGAGGGCCTGTCTCGATACGAACGTTAACCACGTA |
| 7084 | /56-FAM/AGTCGTGTGGGAATCACGACTTCACCATCATTTGTTTTTGCACGACTCCACG |
| 7085 | /56-FAM/TGCGATAGCCCTTTAGGTGACTTTAATAACTGACTCCTTGGACCACGTAGCT |
| 7086 | /56-FAM/CAGACCAGCTGATACCAGTCGTGGTCCGAACGAGACATGCGCGCTAGCCCTT |
| 7087 | /56-FAM/CAGTCGTGTGCACGACTGCCCCTCTAAAGCCTATGCACGACTGTCCGCCCAC |
| 7088 | /56-FAM/TGCGTGTTGTGTGCCCATGTATTCTTTCCACGTAGCTCTCCAGCT |
| 7089 | /56-FAM/GGAGTTTTAAGGTTCACCCGGACTGTTCGG/3BioTEG/ |

[1]Previously published aptamers show the published name and the Maher Lab serial number in parentheses.

[2]Modification abbreviations: /56-FAM/ is an isomer derivative of fluorescein with a six-carbon spacer at the 5' terminus end of the oligonucleotide. + indicates that the following nucleotide caries a locked nucleic acid (LNA) modification. /iSp9/ is an internal triethylene glycol spacer that creates a gap that is not extendable by Taq polymerase during PCR, allowing denaturing electrophoretic gel purification of the top DNA single strand PCR product. /3BioTEG/ is a biotin with an extended triethylene glycol spacer conjugated at the 3' oligonucleotide terminus.
